# Recovery of retinal terminal fields after traumatic brain injury: evidence of collateral sprouting and sexual dimorphism

**DOI:** 10.1101/2025.05.28.656451

**Authors:** Athanasios S. Alexandris, Jaeyoon Yi, Chang Liu, Joseph Belamarich, Zahra Alam, Abhishek Vats, Anthony Peng, Derek S. Welsbie, Donald J. Zack, Vassilis E. Koliatsos

## Abstract

The central nervous system is characterized by its limited regenerative potential, yet striking examples of functional recovery after injury in animal models and humans highlight its capacity for repair. Little is known about repair of pathways/circuits after traumatic brain injury (TBI), which results in disruption of connectivity. Here we utilize a mouse model of diffuse traumatic axonal injury (Impact-acceleration TBI) in order to explore, for the first time, the evolution of structural and functional changes in the terminal fields of the injured visual system. Retinal ganglion cell (RGC) axons and synapses were genetically labeled via AAV transduction, while anterograde and transsynaptic tracers were used to mark terminals and postsynaptic cells. Functional connectivity and visual integrity were assessed by monitoring c-Fos expression following light stimulation and pattern-reversal visual evoked potentials (pVEPs). Our findings demonstrate that, although TAI results in approximately a 50% loss of RGC axons and terminals, surviving RGCs undergo collateral sprouting, a form of compensatory branching of surviving axons, that restores terminal density to pre-injury levels. Transsynaptic tracing and c-Fos mapping confirmed the reestablishment of connectivity, which was also associated with significant improvements in visual function as measured by pVEPs. Interestingly, the recovery process exhibited sexual dimorphism, with female mice showing delayed or incomplete repair. Moreover, collateral sprouting proceeded normally in *Sarm1* knockout mice, evidence of some independence from Wallerian degeneration. Our findings show that collateral sprouting may be an important mechanism of circuit repair in TAI and may represent a promising target for therapeutic interventions.

**Significance:** Homotypic collateral sprouting -the process by which uninjured axons from the same neuronal source extend new branches to reinnervate targets deprived of their original connections- is a fundamental yet understudied mechanism for CNS repair following injury. Unlike heterotypic sprouting, involving sprouting from unrelated pathways, homotypic sprouting offers potential to restore circuit architecture after partial lesions. Here, we employed a model of diffuse axonal injury in the mouse visual system to examine this mechanism. Our research demonstrates surviving retinal ganglion cell axons can re-establish terminal fields, achieving structural and functional connectivity. Importantly, we discovered significant sex differences: female mice showed delayed/incomplete recovery compared to males. These findings provide evidence of repair of brain circuits perturbed by TBI and the role of homotypic sprouting.

## Introduction

Traumatic brain injury (TBI) is associated with a broad range of neuropathologies, including traumatic (diffuse) axonal injury and degeneration, and neuronal cell death (Povlishock, 1992; Stoica and Faden, 2010). Although these pathologies and their functional implications have been extensively investigated in the literature, much less is known about innate processes of regeneration and structural or physiological repair, and the implications of such repair for function. While neural regeneration has been demonstrated in classical examples of neural injury, both in human disease and animal models, it does not always lead to restoration of function (Sulaiman and Gordon, 2013). Specifically in the adult mammalian CNS, restitution of circuits via the precise reconnection of injured neurons with their targets is doubtful (Fawcett, 2020). Instead, the bulk of functional recovery in the CNS is likely achieved by modulation of pre-existing circuits (Makin and Krakauer, 2023) or via the formation of new connections from branching of surviving axons (Cotman et al., 1981; Steward, 1989; Carmichael et al., 2017). The latter is known as collateral sprouting and was first proposed by Exner as a potential mechanism in the peripheral nervoys system (PNS) by which intact nerves reinnervate denervated muscles to prevent atrophy (Exner, 1885). Based on the origin of sprouting axons, collateral sprouting can be defined as homotypic or heterotypic. Homotypic sprouting involves axonal sprouting from surviving neurons within the lesioned pathway and may result in the reinnervation of targets in an anatomically and functionally specific manner. In contrast, heterotypic sprouting is characterized by axonal sprouting from neurons located outside the injured population, While heterotypic collateral sprouting has been extensively studied, especially after severe deafferenting lesions, homotypic collateral sprouting may be the one with the greatest relevance for repair and functional restoration (Cotman et al., 1981; Steward, 1989). The extent and significance of homotypic collateral sprouting in the CNS, however, remains relatively underexplored.

In the present study, we used a well-characterized impact acceleration model of diffuse TBI (IA- TBI) and explored the restoration of terminal fields of the injured visual pathway. The IA-TBI model is characterized by primary injury to axons and ensuing axonopathy termed diffuse or traumatic axonal injury (TAI), in a variety of CNS tracts including the visual pathway (Xu et al., 2016; Ziogas and Koliatsos, 2018; Alexandris et al., 2022, 2023). A unique advantage of the visual system is that its highly compartmentalized anatomy allows the parallel exploration of changes in the retina, optic nerve and tract, and terminal fields in the lateral geniculate nucleus (LGN) and the superior colliculus (SC). We use complementary anatomical and physiological methodologies involving terminal and transsynaptic labeling, c-Fos-based postsynaptic activation, and evoked physiological activity in visual brain areas. To the best of our knowledge, this is the first structural-functional study on the fundamentals of homotypic collateral sprouting after TBI. Our results show evidence of structural and physiological recovery in a time-and sex-specific manner.

## Methods

### Animals

The main study was performed on male and female C57BL/6J mice (Cat. No: 000664, The Jackson Laboratory, Bar Harbor, ME, USA) that were socially housed (2–5 mice per cage) and maintained in a 12-h light/dark cycle, at a temperature of 20–24°C, and with ad libitum access to food and water. For Wallerian degeneration (WD) studies, we used *Sarm1* KO mice backcrossed to the C57BL/6 background bred in our laboratory (Alexandris et al., 2023). *Sarm1* KO mice were generated with homozygous breeding. Congenic C57BL/6 mice were used as controls and referred here as wild type (wt). In *Sarm1* KO mice WD is suppressed (Uccellini et al., 2020; Alexandris et al., 2023).

### Impact acceleration traumatic brain injury in mice

To induce TAI in the visual system (optic nerve), 10- to 14-week old male and female C57BL/6 mice or *Sarm1* KO mice were subjected to IA-TBI or sham injury as described (Alexandris et al., 2023). Briefly, mice were anesthetized with a mixture of isoflurane, oxygen and nitrous oxide, the cranium was exposed, and a 5 mm-thick stainless-steel disc was glued onto the skull midway between bregma and lambda sutures. Then, a 50 g weight was dropped from 85 cm on the metal disk, while the mouse was placed on a foam mattress (4–0 spring constant foam, Foam to Size Inc., Ashland, VA) and its body immobilized with tape. Immediately after injury, the disc was removed and the skull was examined for skull fractures; the rare animals with fractures (<2%) were excluded. The scalp incision was closed with surgical staples. Sham animals underwent the same procedure without the weight drop component. Neurological recovery was assessed by the presence and duration of apnea or irregular breathing and the return of the righting reflex.

Surgical procedures and injuries were performed under aseptic conditions and all animal handling and postoperative procedures were carried according to protocols approved by the Animal Care and Use Committee of the Johns Hopkins Medical Institutions (Protocol Number: M022M442).

### Intravitreal injections

For experiments requiring the use of anterograde or viral tracers, intravitreal injections were performed as follows: Mice were anaesthetised with Ketamine/Xylazine (100 mg/kg, 10mg/kg IP), pupils were dilated with topical tropicamide 1% followed by 0.5% proparacaine HCl for local anesthesia. A hole was made 2 mm posterior to the superior limbus using a sterile 30-gauge (G) needle and 2 μL of virus or tracer were injected into the vitreous via a 33 G needle. Ophthalmic antibiotic gel was applied and mice were allowed to recover. For anterograde tracing experiments, 0.5% of cholera toxin B (CTB) conjugated to Alexa-594 (Cat. No: C22842, Invitrogen, Thermo Fisher Scientific, US) was injected into the left eye 48h prior to perfusion fixation (see below). A period of 48h was allowed for CTB transport based on prior work in rats in which 2 days is sufficient for CTB intensity to reach a plateau (Abbott et al., 2013). To distinguish between technical and biological (between-investigator) variance of the tracing method, in a subcohort of male mice, 0.1% of CTB Alexa-488 (Cat. No: C22841, Invitrogen, Thermo Fisher Scientific, USA) was also injected in the contralateral eye. For axonal and retinotectal synapse labelling, a mixture of AAV2-phSyn1(S)-FLEX-tdTomato-T2A-SypEGFP and AAV2-CMV-Cre were used at a titre of 0.9 x 10^12^ vg/ml each. The plasmid for AAV phSyn1(S)-FLEX-tdTomato-T2A-SypEGFP-WPRE was a gift from Hongkui Zeng (Addgene plasmid # 51509; RRID: Addgene_51509); AAV2 particles were prepared by Vectorbuilder (Chicago, IL, USA). pENN.AAV.CMVs.Pl.Cre.rBG was a gift from James M. Wilson (Addgene viral prep # 105537-AAV2; RRID:Addgene_105537). For transynaptic tracing, AAV2-Y444F-CAG-mWGA-mCherry was used at a titer of 10^13^ vg/ml together with 0.05% CTB Alexa-488; the plasmid was a gift from Xin Duan (Tsai et al., 2022) and AAV2 particles were prepared by Virovek (Houston TX, USA).

### Light stimulation for c-Fos mapping

For the assessment of functional connectivity of the retina with brain targets, injured and sham-injured mice were light-deprived for 60h, exposed to light for 60 minutes, and then returned to darkness for 30 minutes. This schedule was selected based on previous work showing peak c-Fos expression at 90 minutes from stimulus onset (Chaudhuri et al., 2000). Light exposure was performed in a chamber created by lining the walls of a standard mouse cage (20 x 30 x 13cm) with white paper printed with black and white vertical gratings and by attaching a 12 x 1 inch LED strip light fixture producing 1000 Lumen; the top of the cage was then covered with aluminium foil. Animals were exposed to light individually. Mice were stimulated in batches between 9am and 1pm. All animals were perfused with aldehydes at the end of the experiment (see below).

### Electroretinography

Electroretinography (ERG) was performed longitudinally in mice at baseline and various time points after TBI or sham injury using a Celeris system (Diagnosys LLC, Littleton, MA, USA). For each recording session, mice were dark adapted over-night, anesthetized with ketamine/xylazine and recordings were collected using bilateral full field Light Guide electrode-stimulators (Diagnosys LLC, Littleton, MA, USA). Recordings were sampled at 2000 Hz, with each trial lasting 350 ms (including 300 ms post-stimulus). A digital 50/60 Hz notch filter was applied, along with a band-pass filter of 0.125 to 100 Hz. Scotopic ERGs were acquired with flashes of 0.1 cd/m^2^ and 10 s inter-sweep delay. Photopic ERGs were acquired after light adaptation (5 minutes at 9 cd/m2) and with flashes of 3 cd/m2. Flicker ERGs were acquired using 10 Hz flashes of the same intensity. Extracted values (*a*-wave, *b*-wave and N1-P1 amplitudes) were then averaged across individual trials and eyes per mouse.

### Pattern visual evoked potentials: recordings and analysis

Mice were subjected to IA-TBI or sham injury and after a brief recovery period (10 minutes), subdural electrodes were implanted over the right visual cortex (V1; AP: -3.0 mm, ML: -1.7 mm) and frontal cortex (AP: 1.7 mm, ML: 1.7 mm). On the day of recording, mice were anesthetized with ketamine/xylazine, and pattern-reversal visual evoked potential (VEP) recordings were conducted using visual pattern stimulation (CELERIS, Diagnosys LLC). Recordings were sampled at 2000 Hz, with each trial lasting 700 ms (including 514 ms post-stimulus). A digital 50/60 Hz notch filter was applied, along with a band-pass filter of 1 to 100 Hz for VEP signals. No baseline or drift correction was applied. The visual stimuli consisted of vertically oriented gratings of different contrast and spatial frequency combinations (100/0.125, 100/0.25, 100/0.31, 100/0.38, 70/0.125, 70/0.25, 70/0.31, 80/0.38, 40/0.125, 50/0.25, 50/0.31, 60/0.38, 40/0.25, 45/0.31) that reversed at 1.4 Hz and were presented with a mean luminance of 150 cd/m^2^. Each of the 14 stimuli was presented sequentially, with 15-40 trials recorded for each stimulus. After all stimuli were tested, this sequence was repeated 7 to 8 additional times (∼1h per animal). Time-locked average responses were used to estimate the P1-N1 amplitude of the pVEP wave.

The relationship between response amplitude and stimulus parameters was modelled using a generalized additive model (GAM), implemented in the mgcv package (Wood, 2011) in R (v4.1.2) and was specified as:

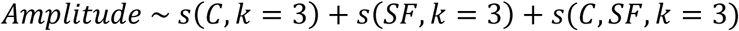

where 𝑆(·) represents smooth functions modeling the effect of contrast, C, spatial frequency, SF, and their interaction. The smoothing parameters were estimated using restricted maximum likelihood to balance flexibility and prevent overfitting. Separate models were fit for each group to allow for group-specific differences in response patterns and were illustrated as heatmaps based on a global color-scale whose range is derived from the 5th and 95th percentiles of all raw amplitude values. For each experimental group, sex, and time point, a threshold contrast curve was derived to quantify the minimum contrast required to elicit a VEP response at each spatial frequency. The threshold was defined as the lowest contrast at which the GAM-predicted amplitude exceeded 50% of the maximum amplitude observed in the corresponding sham cohort. This threshold was chosen to ensure that the derived values stayed within the range of the observed measurements. To smoothen variability in the raw threshold estimates, locally weighted scatterplot smoothing (LOWESS) was applied with a span of 0.5. To derive bias-adjusted threshold contrast curves and quantify uncertainty, we used a leave-one-out jack-knife resampling procedure (Efron and Stein, 1981), iteratively removing each subject from the dataset, refitting the GAM, and recomputing the threshold contrast curve. The resulting variability was used to adjust average threshold values and estimate standard errors for plotting and statistical comparisons.

### Tissue processing and immunohistochemistry

Mice were anesthetized with 7.5% chloral hydrate (Sigma, C8383-1KG) in normal saline and transcardially perfused with 0.1 M PBS for 1-2 minutes and then with freshly depolymerized 4% paraformaldehyde in PBS (pH 7.4) Brains were postfixed overnight in the same fixative at 4 °C with mild shaking. Optic nerves, brains and retinas were dissected and cryoprotected in 20% glycerol/5% DMSO. Brains were sectioned on a freezing microtome (40 μm) through the tectum and sections were stored in antifreeze solution (50% 0.1M phosphate buffer, 30% sucrose, 1% polyvinylpyrrolidone, 30% ethylene glycol in ddH2O) at -20 °C until staining.

#### Immunohistochemistry-brain sections

For the analysis of retinal terminal fields in SC and LGN, serial floating sections (1:6) were immunostained, mounted on slides, air dried, and coverslipped with Vectashield (Vector Laboratories Inc., H-1000-10). For immunostaining, brain sections were washed in 0.1M tris buffered saline (TBS; 3 x 30 mins) and then blocked and permeabilized with 5% normal goat serum and 0.4% Triton X-100 in TBS for an hour at room temperature under agitation. Sections were then incubated in primary antibodies (**Table 1**) diluted in 5% normal goat serum, 0.1% Triton-X and 0.1% NaN_3_ in TBS mild shaking overnight at 37 °C. After rinsing in TBS, sections were incubated in corresponding secondary antibodies (Alexa-Fluor conjugated antibodies, 1:300) in the same diluent overnight at 37 °C. Sections were then counterstained with DAPI washed before mounting, air drying and coverslipping as before.

**Table 1.** Primary antibodies used in this study.

| Primary Antibody Target, Host | Product No | Dilution Ratio, target sample | Incubation Time | Incubation Temp |
| --- | --- | --- | --- | --- |
| Neuronal Nuclear antigen (NeuN), Guinea Pig | ABN90P, Millipore Corp | 1:1000, brain section | Overnight | 37 °C |
| c-Fos, Rabbit | 226 008, Synaptic Systems, | 1:2000, brain section | Overnight | 37 °C |
| Red fluorescent protein (RFP), Rabbit | 600-401-379, Rockland Immunochemicals, | 1:1000, brain section | Overnight | 37 °C |
| Green fluorescent protein (GFP), Chicken | A10262, Invitrogen, | 1:500, brain section | Overnight | 37 °C |
| Myelin basic protein (MBP), chicken | C-1383-50, Biosensis | 1:500, brain section | Overnight | 37 °C |
| Gamma synuclein (SNCG), mouse | 10579-316, Abnova | 1:500, whole retinas | 36 hours | 4 °C |
| Beta-Tubulin III, TUJ1, rabbit | Cat# 802001, BioLegend, | 1:100, ultrathin ON sections | Overnight | 4 °C |

#### Immunohistochemistry-optic nerve ultrathin sections

In order to count axons in optic nerves accurately and reliably, we developed a high-resolution IHC procedure for ultrathin sections (75 nm) from resin-embedded optic nerves that affords single-axon resolution (Liu et al., 2025). In brief, optic nerves were washed =in 3.5% sucrose and 50 mM glycine in 0.01 M PBS at 4 °C under agitation, dehydrated in ethanol series, and embedded in LR White resin (Cat# 14381, Electron Microscopy Sciences, Hatfield, PA).Resin-embedded ONs were sectioned into 70 nm ultrathin sections. Sections from the pre-chiasmatic, distal ON segment were blocked and permeabilized with 5% normal goat serum and 0.5% Triton X-100 in PBS for 1 hour at room temperature in a humidifying chamber. Sections were then incubated with a TUJ1 rabbit antibody (see table 1) in blocking buffer overnight at 4 °C. After rinsing, sections were incubated with an Alexa-594- conjugated goat anti-rabbit in blocking buffer for 2 hours at room temperature. Sections were then rinsed and counterstained with DAPI. Sections were dried and mounted with a Vectashield® anti- fade medium.

#### Immunohistochemistry-retinas

Retinas were dissected and washed in PBS, and then permeabilized and blocked with 0.3% Triton X-100 and 10% normal goat serum n PBS for 2 hours at room temperature. Retinas were then incubated with a SNCG mouse antibody in blocking buffer for approximately 60 hours at 4 °C under agitation. Subsequently, retinas were incubated with Alexa-647-conjugated goat anti-mouse IgG in blocking buffer for approximately 36 hours at room temperature, washed in PBST (x3) and flat mounted with Prolong Diamond Antifade Mountant (Cat. No: P36970, Invitrogen, Thermo Fisher Scientific, Waltham, MA).

In all experiments tissues from sham and TBI groups were processed in the same batches.

### Microscopy and image analysis

#### Quantitative analysis of terminal fields

For morphometric analysis of retinal terminal fields in the SC and LGN, confocal images of serial sections (1:6) were acquired with a MICA confocal microscope (Leica Microsystems, Deerfield, IL) at 63x magnification and tilling. Single optical sections were acquired at a distance of 10 µm from the edge of the section using consistent acquisition settings. Tilled images were imported to FIJI (NIH, MD, USA)(Schindelin et al., 2012) and transformed by Gaussian blurring (sigma = 2) before global thresholding with a fixed threshold for each signal for the generation of binary maps. The percent area coverage of each signal (CTB, tdTomato or SypGFP), was then analyzed in each region of interest by the “Analyze Particles” function. The mean value of all images per region and per case was calculated. To assess SC atrophy, confocal images of adjacent MBP-stained sections were used to trace the superficial layer of the SC at 20x.

#### Estimates of c-Fos(+) and WGA(+) neuron ratios

Images were acquired using a confocal microscope as before under 20x magnification. Select regions of interest (ROIs) from tiled images were cropped and imported to the commercially available AI platform Biodock (Biodock, AI Software Platform, 2024), and an analysis model was trained manually for segmentation of NeuN(+) neurons and calculation of intensity values per neuron. Raw data including individual intensity measurements per channel for all segmented NeuN(+) neurons, were analyzed to calculate the ratio of NeuN(+) neurons that express c-Fos or WGA. Specifically, for the c-Fos experiment, a minimum intensity threshold was determined by comparing the inverse cumulative distribution of c-Fos labeling intensities between unstained, non-stimulated and light- stimulated animals and by selecting the minimum intensity threshold that separates the two groups based on non-overlap of 95% CIs (**Fig. S1**). Then the ratio of NeuN(+) and c-Fos(+) neurons was determined per case and per brain region. The ratio of WGA(+) positive neurons was similarly calculated with the exception that the experiment was performed in two separate batches that resulted in different raw intensities of WGA signals. Therefore we defined the threshold for positive WGA(+) signal in each group as the average value of the median WGA intensity drawn from each sham animal. In addition, because we were not able to estimate the % of RGCs transduced due to very low level expression of WGA in retinas, we excluded cases in which the fraction of WGA(+) neurons in the SC was below 5%.

#### RGC counts

Confocal images of flat-mounted retinas immunostained with SNCG were acquired under 20x magnification. Twelve images were acquired per retina covering the central, middle, and peripheral regions of each quadrant. The Biodock platform was used to estimate numbers of SNCG-labeled retinal ganglion cells as before (Liu et al., 2025).

#### Axon counts

Confocal images of ultrathin stained sections were acquired at 60x magnification and tilling. Three sections per ON were imaged for quantitative analysis using FIJI. Axons were segmented using the Watershed plugin, and individual axons were counted using the “Analyze Particles” function. A mean axon count was then calculated for each ON section.

All quantifications were performed by an experimenter blind to group designation.

### Statistical analyses

Graphpad Prism 9.1 (GraphPad Software, La Jolla, Ca, USA) and R Studio (2022.2.0.443) was used for statistical analyses and plotting of figures. Details of statistical analyses are presented in figures and **Supplemental statistical analyses (Tables S1-S9)**. Significance threshold (*p*) was set at 0.05. Unless otherwise specified, multiple comparisons were corrected using the Holm– Šídák method.

## Results

### 1. Impact Acceleration TBI leads to RGC perikaryal, axonal and terminal degeneration followed by late reconstitution of terminal fields

We have previously shown that a single IA-TBI event in the adult mice generates a well-defined injury front within the pre-chiasmatic ON that results in partial disconnection and subsequent slow degeneration of 40% of axons as well as perikaryal degeneration of 20% of RGCs (Alexandris et al., 2023). To map and quantify changes in RGC terminals, we performed unilateral intravitreal injections with CTB conjugated to Alexa-594 48 hours before termination of experiment at 7, 14, 28 or 56 days after IA-TBI or sham-injury in male and female mice (**Fig. 1 A-B**). Mouse RGCs project to more than 50 retinorecipient regions, with the contralateral SC and LGN being the major targets (Martersteck et al., 2017). Because approximately 90% of RGCs (including the majority of LGN projecting RGCs) project to the SC (Ellis et al., 2016), we focused our analyses on changes in the density of retinocollicular terminals in the superficial layer of the contralateral SC.

**Fig. 1.**
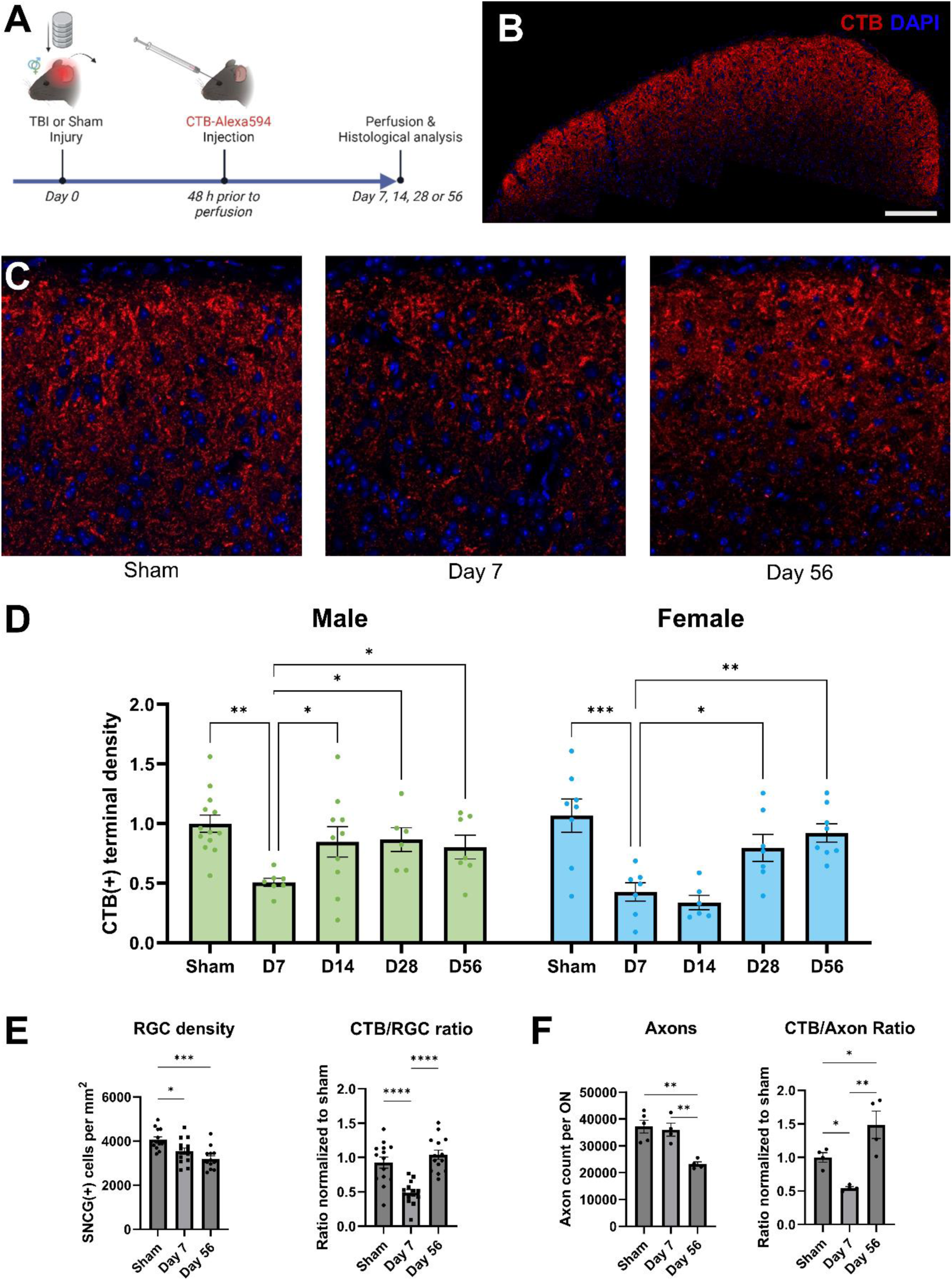
Course of collateral sprouting in retinocollicular projections after IA-TBI in male and female mice. **A.** Experimental design. **B-C.** Confocal images showing retinocollicular projections traced by intravitreal CTB injections (red). Panel (C) shows representative images of the retinorecipient superficial layer of the superior colliculus in sham and injured male mice (C). **D.** Changes in CTB(+) terminal density after IA-TBI. In both male (green) and female (blue) mice there is ∼50% loss of CTB(+) terminals at 7 days followed by recovery to pre-injury levels in a sexually dimorphic manner. **E-F**. Normalization of retinocollicular terminal densities to RGC (E) and axon counts (F). Left panels shows densities of γ-synuclein(+) RGCs in retinas (E) or TuJ1(+) axon counts in optic nerves (F) from pooled male and female mice. Right panels show terminal densities per RGC or axon. For details see **supplemental statistical analyses** Table S1.

As expected from the magnitude of axon loss following IA-TBI, we found that the density of CTB(+) terminals was reduced by more than 50% at 7 days post-injury in both male and female mice. This loss of terminals was followed by nearly complete recovery to pre-injury levels but in a sexually dimorphic manner. In males, the density of terminals recovered by 14 days post-injury, whereas female mice had a significantly slower course, with evidence of recovery at 4 weeks after injury (**Fig. 1C-D; Table S1**). There was no significant difference between male and female mice at baseline (t(19.94)=0.46, *p*=0.6). To exclude that the observed increase in CTB(+) terminals was confounded by atrophy of the superficial layer, we measured the surface area of the superficial layer of the SC based on MBP immunohistochemistry (Byun et al., 2016), and we found no significant atrophy and no effect on CTB(+) terminal measures (**Fig. 2**). In view of the fact that there is no axonal regeneration in the ON after IA-TBI (Alexandris et al., 2023), the above findings suggest that the increased density of retinocollicular terminals after 7 days is due to collateral sprouting of surviving axons. Indeed, if we divide the density of CTB(+) terminals by the number of RGC perikarya or axons we estimate that, on average, the projections of each surviving RGC increase nearly by two-fold (**Fig. 1E-F;Table S1**).

**Fig. 2.**
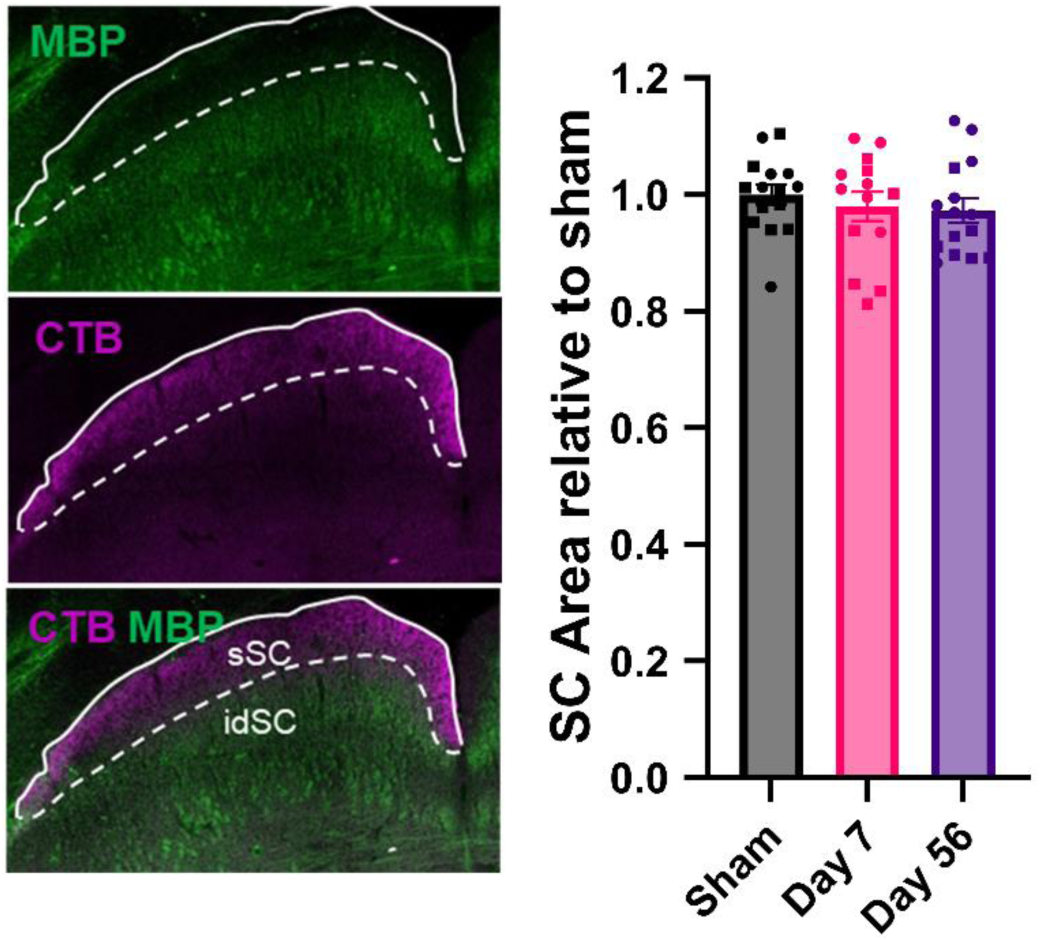
IA-TBI is not associated with atrophy of the superficial layer of the superior colliculus (sSC). Total area of the superficial layer was measured in 1:6 series based on MBP immunohistochemistry. (Left) Confocal image of a section through the superior colliculus showing definition of the superficial layer based on MBP immunoreactivity (green) with white interrupted line. (Right) Comparison of surface areas between groups in pooled male and female mice. There is no change in after IA-TBI (F (2, 41) = 0.47; *p*=0.63; Sham vs. Day 7: t(41)=0.69; *p*=0.58; Sham vs. Day 56: t(41)=0.94; *p*=0.58).

We also noted that the density of retinocollicular projections exhibit substantial variance among individual mice, including subjects in the sham-injured group (coefficient of variation of 30%).

Therefore, to distinguish whether the source of variance is biological or technical, we assessed an independent group of injured and sham-injured mice (28 days after injury) with additional intravitreal injection of CTB Alexa-488 in the contralateral eye. We found that measures of retinocollicular terminal densities were highly correlated between left and right eyes (Pearson’s r = 0.86, *p*<0.0001) while nested variance component analysis revealed that inter-individual differences accounted for more than 50% of the observed variance (5-fold of that attributed to between-eye differences) (**Fig. 3**; **Table 2**). Therefore, we surmise that the observed variance in the density of retinocollicular projections stems from biological rather than technical factors.

**Fig. 3.**
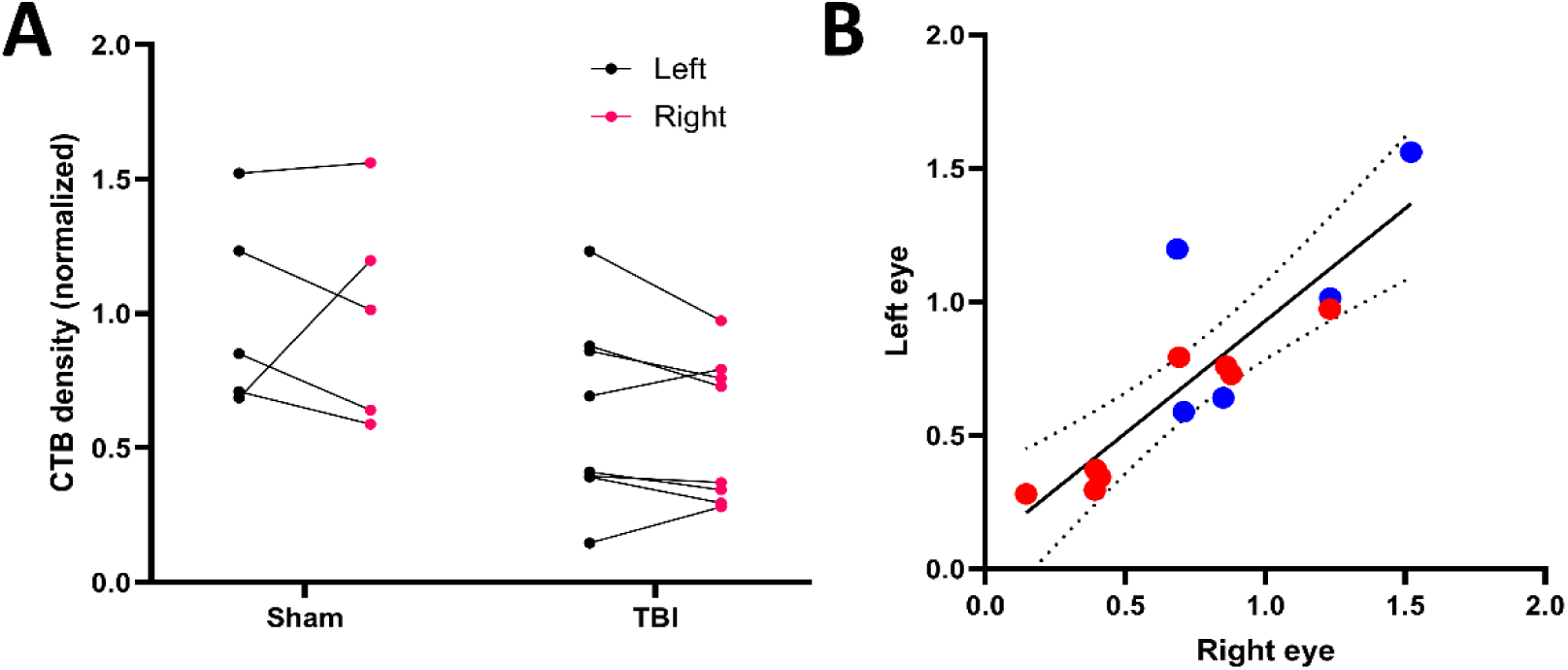
Assessment of intra and inter-individual variance in retinocollicular projections. Injured or sham male mice received intravitreal injections of CTB-Alexa-488 and CTB-Alexa-594 in the right and left eyes, respectively, before perfusion at 28 days following injury. **A-B** CTB densities in corresponding superior colliculi show significant correlation with Pearson *r*= 0.86, *p*<0.0001. In panel (B) sham animals are represented by blue circles, TBI animals are represented by red circles.

**Table 2.**
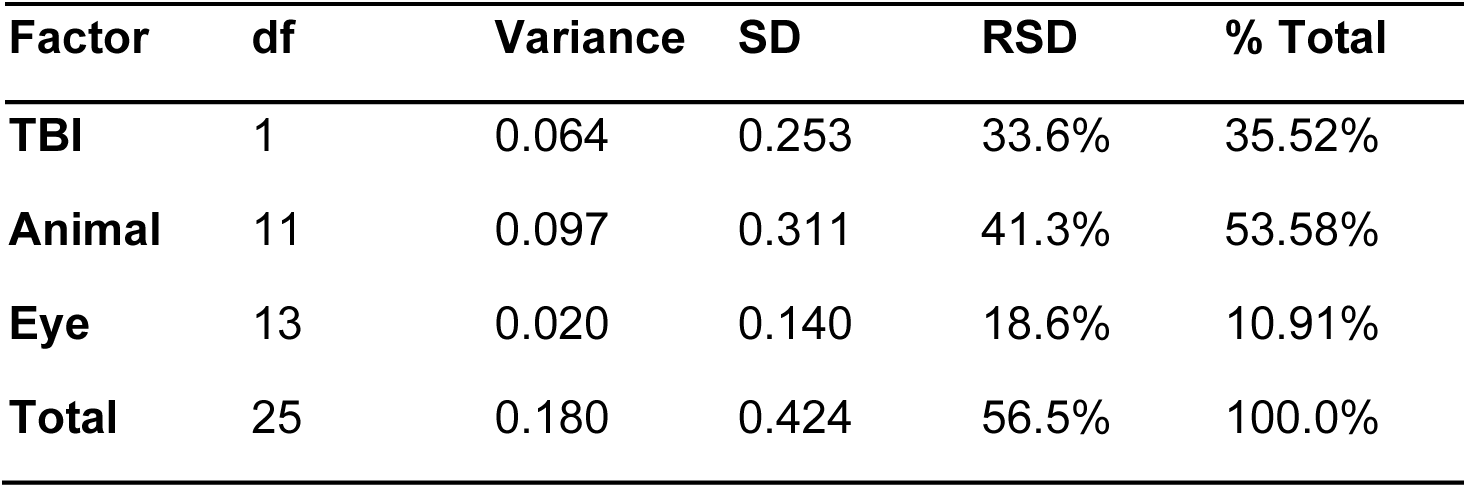
Results of variance component analysis. df, degrees of freedom; SD, standard deviation; RSD, relative standard deviation. See text for discussion.

| **Factor** | **df** | **Variance** | **SD** | **RSD** | **% Total** |
| --- | --- | --- | --- | --- | --- |
| **TBI** | 1 | 0.064 | 0.253 | 33.6% | 35.52% |
| **Animal** | 11 | 0.097 | 0.311 | 41.3% | 53.58% |
| **Eye** | 13 | 0.020 | 0.140 | 18.6% | 10.91% |
| **Total** | 25 | 0.180 | 0.424 | 56.5% | 100.0% |

Finally, because CTB labeling depends on anterograde transport which could be affected by the course of axonopathy following IA-TBI, we further confirmed changes in retinocollicular terminals in a small cohort of male mice whose projections were labeled prior to injury with a genetic approach using AAV-mediated expression of cytosolic tdTomato and synaptophysin fused to GFP (sypGFP) for marking axons and synapses, respectively (**Fig. 4A-C**). Pre-injury labeling with CTB was not feasible due to its limited persistence (∼2 weeks), whereas AAV-based labeling offers stable, long-term expression. As we observed with the CTB-based experiment (Fig.1), we found that the density of both terminal axons (tdTomato(+)) and synapses (sypGFP(+)), decreased by 50% 7 days after injury and were restored to pre-injury levels by day 28 (**Fig. 4D; Table S2**).

**Fig. 4.**
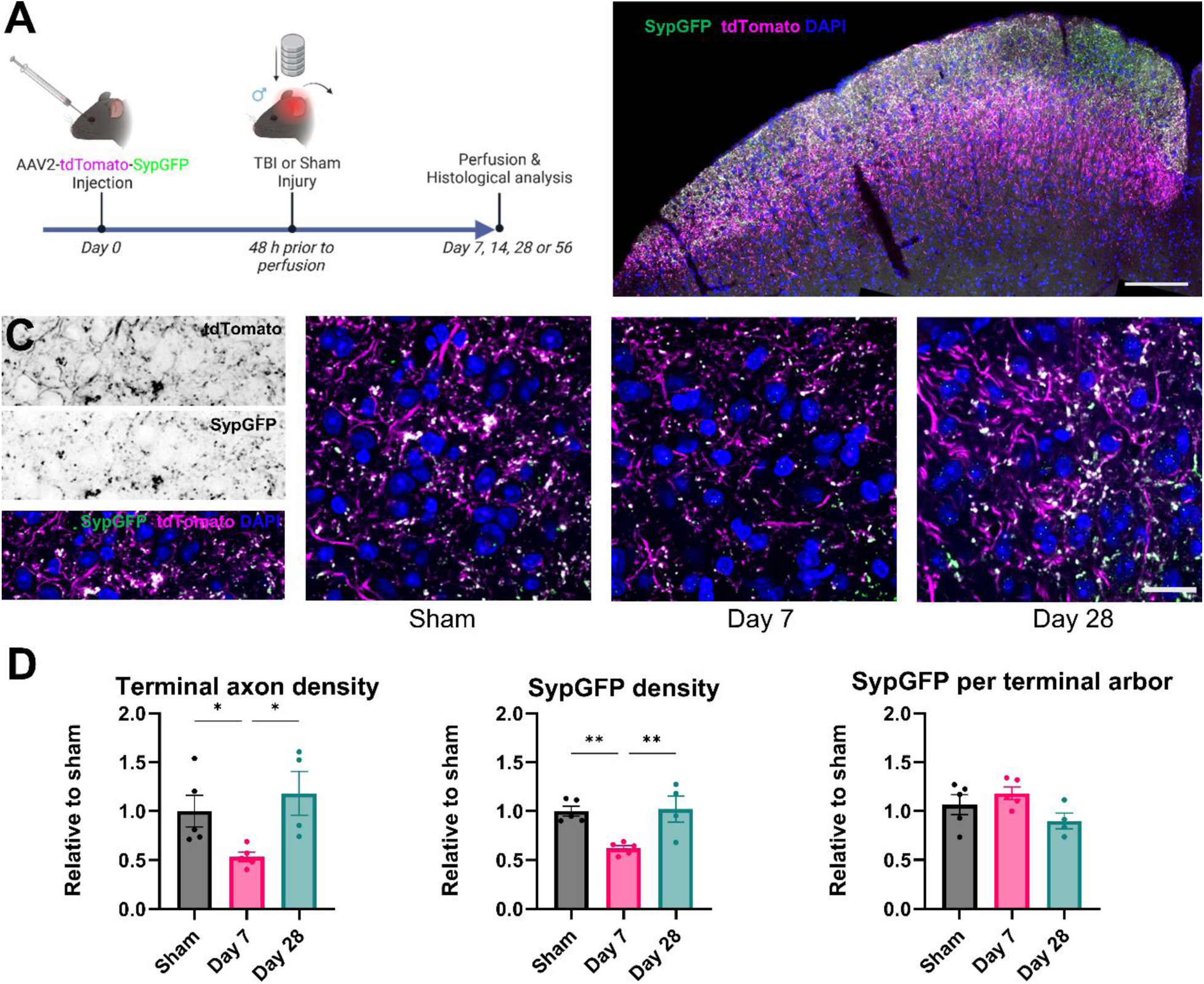
Course of sprouting in retinocollicular projections after IA-TBI using genetic tracers. **A.** Experimental design. **B-C.** Confocal depiction of traced retinocollicular projections. Terminal axons (tdTomato(+) are shown in magenta and synaptic terminals (SypGFP) in green. **D.** Changes in densities of terminal axons based on tdTomato labelling (left panel), synaptic terminals based on SypGFP (middle panel), and ratio of synapses per terminal axon arbors (right panel). IA-TBI leads to equal loss of synapses and terminal axons which is followed by their reconstitution by 28 days. For details see **supplemental statistical analyses** Table S2.

### 2. Terminal field recovery and re-establishment of functional connectivity

While previous results provide strong evidence of structural plasticity and the regeneration of retinocollicular terminals to pre-injury levels within two to four weeks post-injury, it remains unclear whether the new collaterals form mature or functional synapses with target neurons at levels approximating pre-injury baselines. To address this question we first used the genetically encoded transynaptic tracer wheat germ agglutinin-fused to mCherry (WGA-mCherry)(Tsai et al., 2022).

Wheat germ agglutinin is transported across synapses and mCherry is resistant to degradation and can thus accumulate in lysosomes (Tsai et al., 2022). Therefore the extent of WGA-mCherry accumulation in post-synaptic neurons serves as a surrogate of SC connectivity via mature synapses. In this experiment, injured or sham-injured mice were unilaterally injected with AAV- WGA-mCherry and CTB-Alexa 488 7 days prior to perfusion at 7, 14 and 56 days following injury (**Fig. 5A-C**). We then assessed the fraction of NeuN(+) neurons which expressed WGA-mCherry. In an effort to increase statistical power given the substantial technical variability associated with viral vector delivery in this experiment, data from both sexes were pooled for analysis. This analysis revealed that IA-TBI causes a significant loss of postsynaptic labeling, subsequently followed by a return to pre-injury levels (**Fig. 5D; Table S3**), and recovery of postsynaptic connectivity. As was the case in our CTB and synGFP experiments, female mice showed a trend toward delayed recovery.

**Fig. 5.**
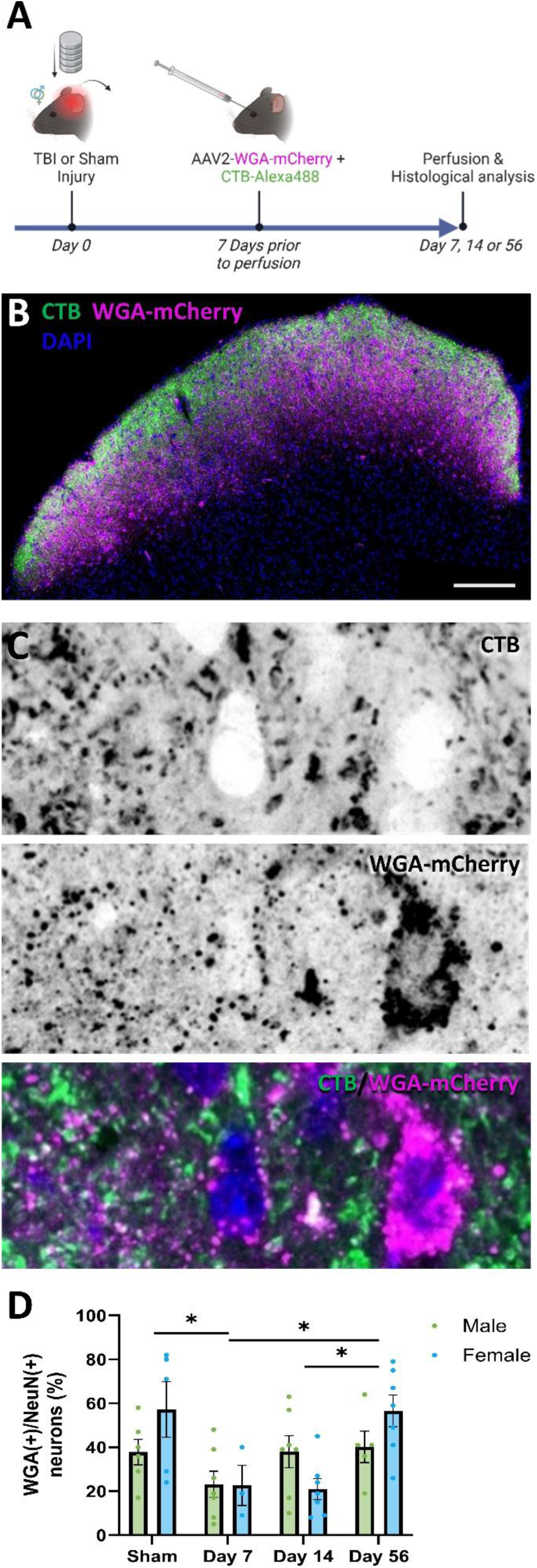
Transsynaptic tracing of retinocollicular connectivity after IA-TBI. **A.** Experimental design. **B-C.** Confocal depiction of retinocollicular projections (green) and transsynaptically labelled post- synaptic neurons with WGA-mCherry (magenta). **D.** Changes in retinocollicular connectivity based on WGA-mCherry labeling of Neun(+) superior collicular neurons in male (green) and female (blue) mice. In the first week after injury, there is significant loss of connectivity followed by gradual recovery. For post-hoc comparisons male and female mice have been pooled – For details see **supplemental statistical analyses** Table S3.

In order to further assess the functionality of newly formed mature synapses, we performed c-Fos labeling in a paradigm of light stimulation after a brief period of light deprivation (**Fig. 6A-B**).

**Fig. 6.**
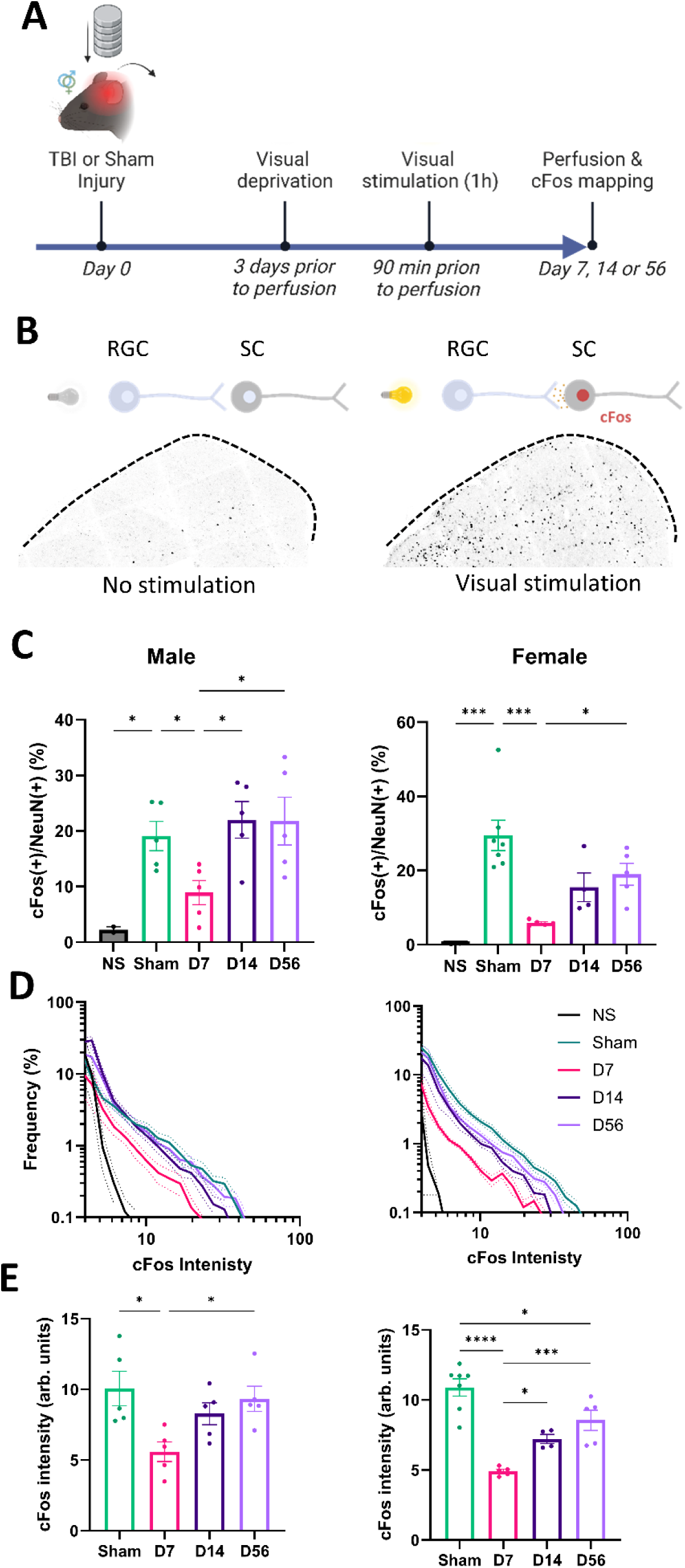
c-Fos based analysis of retinocollicular connectivity after IA-TBI. **A.** Experimental design. **B.** c-Fos immunoreactivity in the superior colliculi of control mice and mice exposed to visual stimuli (see diagram on top). **C.** Changes in retinocollicular connectivity based on c-Fos expression in Neun(+) SC neurons in male and female mice. In the first week post-injury, there is significant loss of connectivity followed by recovery by 14 days in male mice and 56 days in female mice. **D.** Relative frequency distribution plots for c-Fos intensities per Neun(+) neuron. **E**. Median c-Fos intensities in the top 10% of neurons. In contrast to c-Fos(+) ratios, c-Fos intensities show a delayed recovery in both male and female mice, and only partial recovery in female mice. For details see **supplemental statistical analyses** Table S4.

Expression of c-Fos is associated with neuronal activation and correlates with specific patterns of neuronal activity, including synchronized firing and burst activity (Guo et al., 2007; Yassin et al., 2010; Anisimova et al., 2023). While not a direct measure of firing frequency, c-Fos expression in retinorecipient areas in response to light exposure can serve as a marker of functional synaptic transmission and network-level activation. In sham-injured animals, light stimulation leads to expression of c-Fos in 24 and 36% of NeuN(+) cells in the SC of male and female mice, respectively, compared to ∼ 1.8% at baseline. Seven days after injury the fraction of c-Fos responsive neurons decreased by 50 and 80% in male and female mice, concordant with loss of retinocollicular connectivity. This was, again, followed by recovery to pre-injury levels over the duration of the experiment but in a sexually dimorphic manner, with females recovering more slowly than males (**Fig. 6C; Table S4**).

A binary assessment of c-Fos expression may mask the nuances of neuronal activation levels. By plotting a frequency distribution of c-Fos intensities, we found that overall c-Fos expression follows a lognormal or power law distribution, where high c-Fos levels are encountered in only a very small fraction of neurons (**Fig. 6D**). This distribution is consistent with previously described network dynamics, whereby the majority of neurons fire at low rates and only a minority (∼10%) of neurons exhibit high-frequency firing patterns, often associated with critical information processing or heightened synaptic activity (Buzsaki and Mizuseki, 2014) . Therefore, to further assess whether the recovery of retinocollicular connectivity is accompanied by corresponding functional activation at the network level, we evaluated changes in the median c-Fos intensity of the top 10% of neurons. While not directly measuring firing frequency, this approach allows us to identify the subset of neurons most responsive to coordinated circuit activity in response to light stimulation. In contrast to the overall recovery of connectivity, we found that the activity of the top 10% of neurons recovered much more slowly, and in female mice it did not return to baseline levels even two months after injury (**Fig. 6E; Table S4**).

To better understand the impact of TAI upstream of the main visual pathway, we also examined retinogeniculate connectivity in the LGN using CTB labeling, c-Fos mapping, and WGA-mCherry tracing. For these analyses, data showed higher variability and data from both sexes were pooled to increase statistical power, with results showing trends similar to the effects of IA-TBI on retinocollicular projections (see **Fig. 7; Table S5**).

**Fig. 7.**
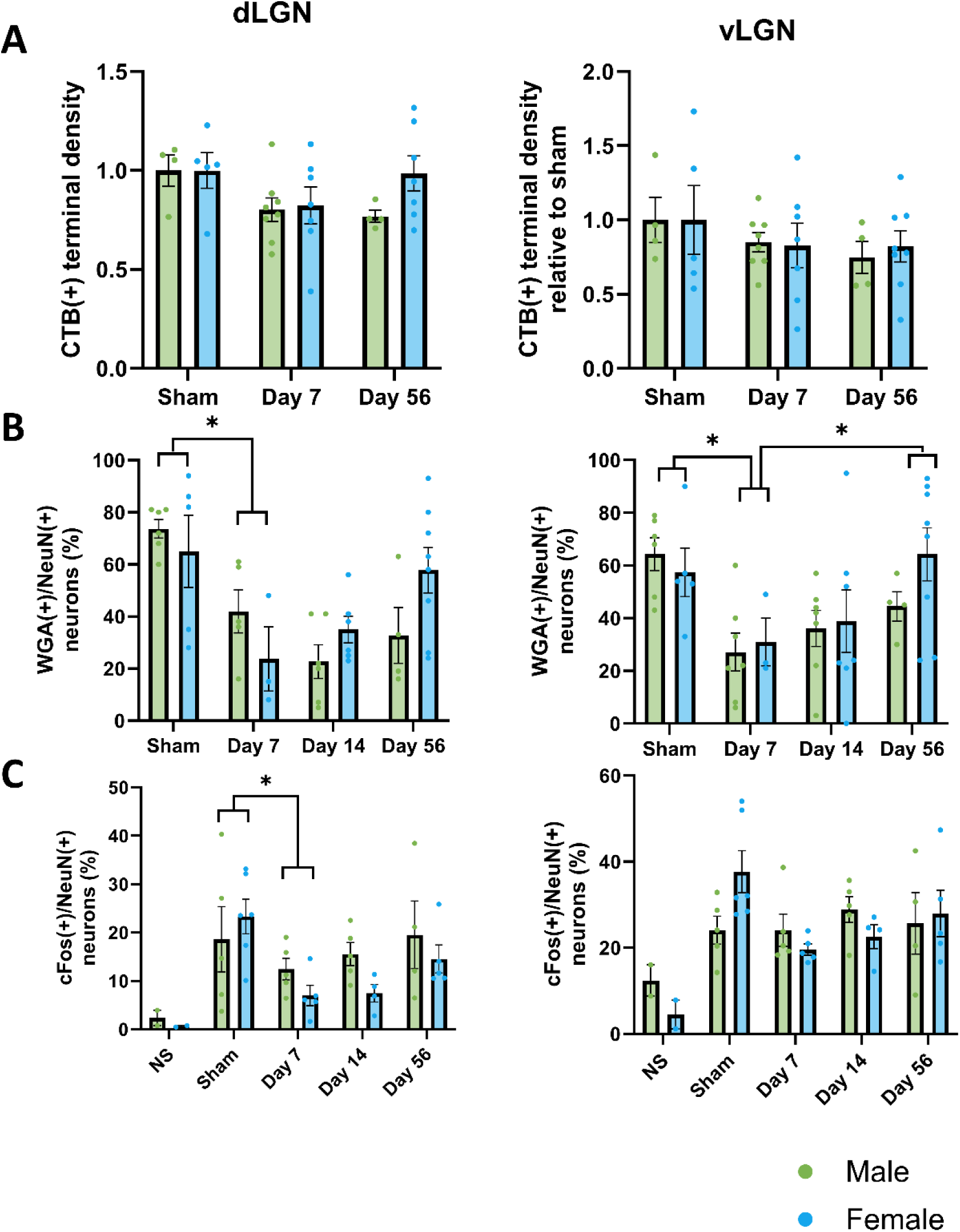
Analyses of retinogeniculate sprouting and connectivity. **A.** CTB(+) terminal densities of the contralateral dorsal and ventral lateral geniculate nucleus (dLG and vLG) as per Fig.1. **B.** Transynaptic-tracing based connectivity analysis in the dLG and vLG as per Fig.4. **C.** c-Fos based connectivity analysis in the dLG and vLG as per Fig. 5. For details see **supplemental statistical analyses** Table S5.

### 3. Changes in pattern visual evoked responses in the course of collateral sprouting

To evaluate the functional consequences of IA-TBI on the visual system, we initially tested standard approaches used to assess visual pathway integrity. Optomotor response testing did not reveal consistent deficits in this model, and flash visual evoked potentials (VEPs) likewise failed to show measurable alterations in cortical responsiveness (unpublished observations). Because both methods lack sensitivity for detecting subtle changes in visual function, we turned to pattern VEPs, which provide a more robust measure of contrast sensitivity and visual acuity thresholds.

To exclude direct injury to the retina, we performed electroretinography (ERG) for the assessment of photoreceptor and bipolar cell function. Despite some inter-session variation in amplitudes, scotopic, photopic and flicker ERGs indicated no significant deficits in retinal responses to full-field stimulation (**Fig. 8; Table S6)** consistent with preserved retinal function after injury.

**Fig. 8.**
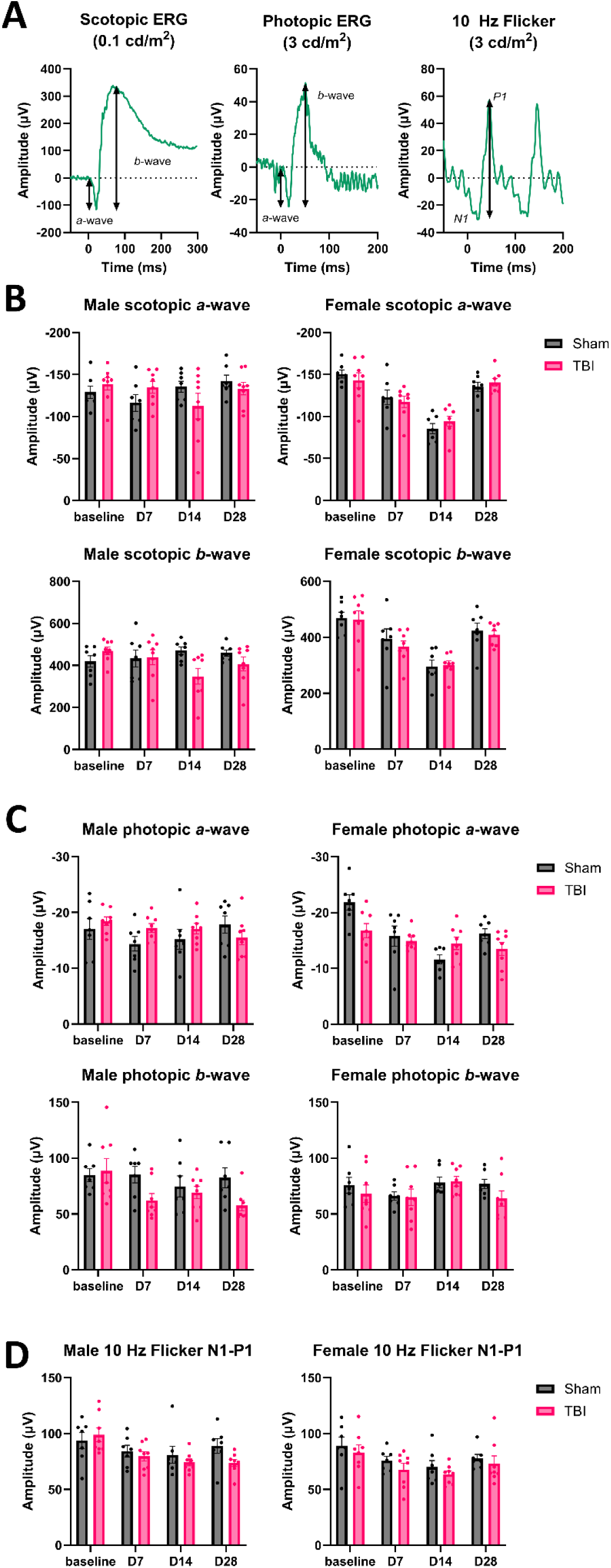
Full field electroretinography (ERG) shows no deficits in retinal function after IA-TBI in male and female mice. **A**. Representative waveforms for scotopic, photopic and flicker flash ERGs. **B.** Scotopic *a*- and *b*-wave amplitudes. **C**. Photopic *a*- and *b*-wave amplitudes. D. 10 Hz Flicker N1- P1 amplitudes. There are no significant differences between TBI and sham mice. For details see **supplemental statistical analyses** Table S6.

To examine the impact of compromised connectivity of retinal efferents, we developed a protocol using pattern reversal VEPs using gratings of varying spatial frequencies and contrasts (**Fig.9A-C**). This approach enables us to estimate the lowest contrast at which the cortical response exceeded a predefined detection threshold—a physiological proxy for the contrast sensitivity function. Heatmaps of absolute pVEP amplitudes in sham animals demonstrated a consistent, time-dependent increase in maximum response amplitudes (**Fig. 9 D-E**; **Fig. S2 A; Tables S7-8**). To account for these changes, we defined relative contrast thresholds based on sham responses (see Methods) and compared threshold curves between TBI and sham groups at each time point (**Fig. 10**). In both sexes, IA-TBI narrowed the functional contrast range, an effect signifying that, for each spatial frequency, higher contrast levels are required to elicit a cortical response comparable to the sham scenario. These deficits seem to recover over time in a sex-dependent manner: males returned to sham-like thresholds by day 14, whereas females exhibit persistent deficits, as indicated by the sustained divergence of their threshold curves. A similar pattern is seen when an absolute global threshold is used instead, with the key difference that female mice show partial recovery at 56 days (**Fig. S2 B-C**).

**Fig. 9.**
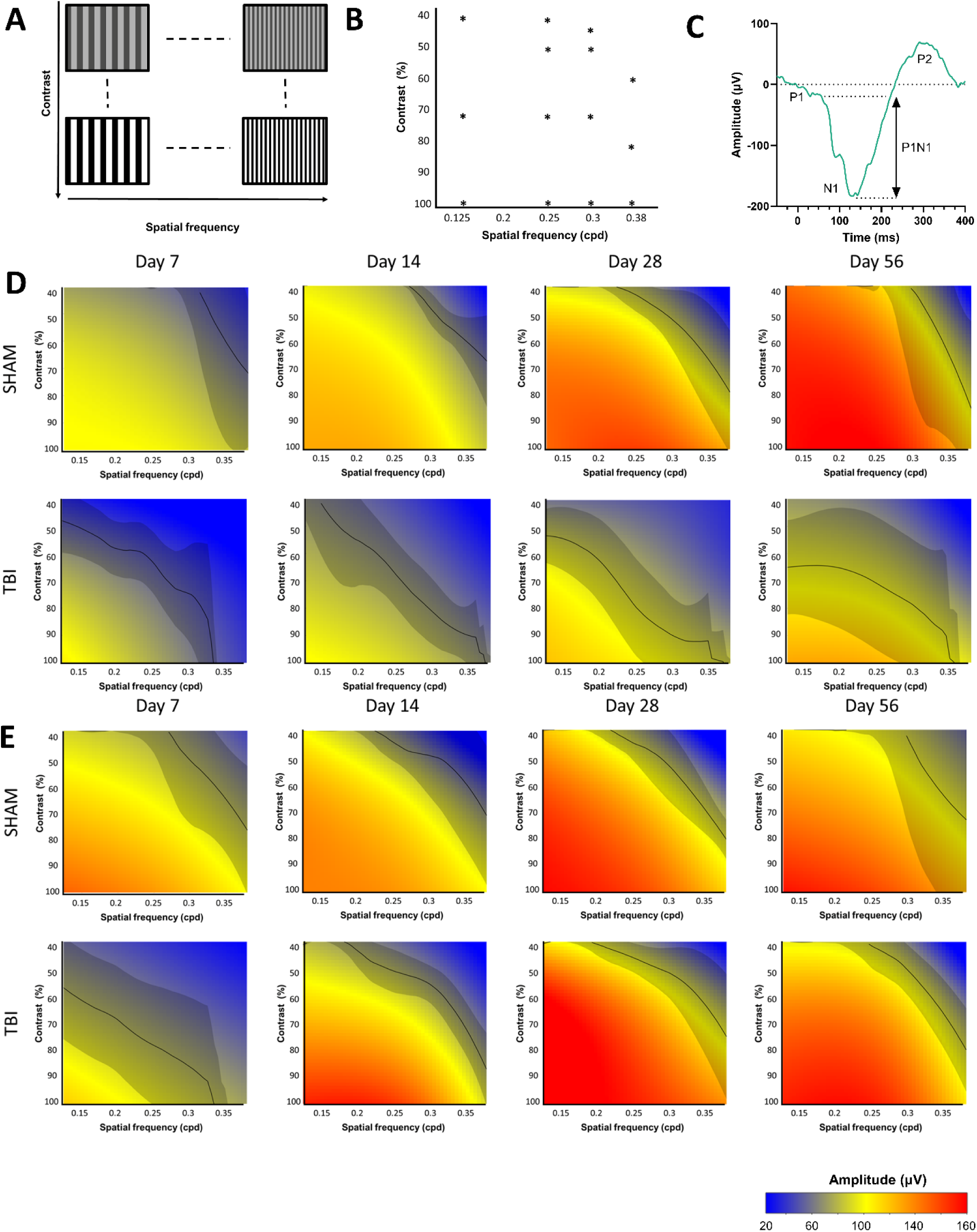
Contrast sensitivity assessment based on pattern reversal visual evoked potentials (pVEP). **A-B**. Range of black and white gratings of different spatial frequency and contrast used for pVEPs. **C.** Example of stimulus-locked averaged tracing of pVEP showing typical P1-N1-P2 waveform. **D-E.** General additive model-based heatmaps of pVEP amplitudes in response to stimuli of different spatial frequency and contrast combinations for female (D) and male (E) mice. Heatmaps are colorized based on a global scale from the bottom and top 95^th^ percentiles for all cases and time-points assessed. In (D) and (E), black-lines indicate bias-adjusted contrast sensitivity thresholds with 95% confidence intervals (shaded gray) derived from the 50% of the maximum amplitude for the sham group per sex and time-point. For details of model parameters see **supplemental statistical analyses** Table S7.

**Fig. 10.**
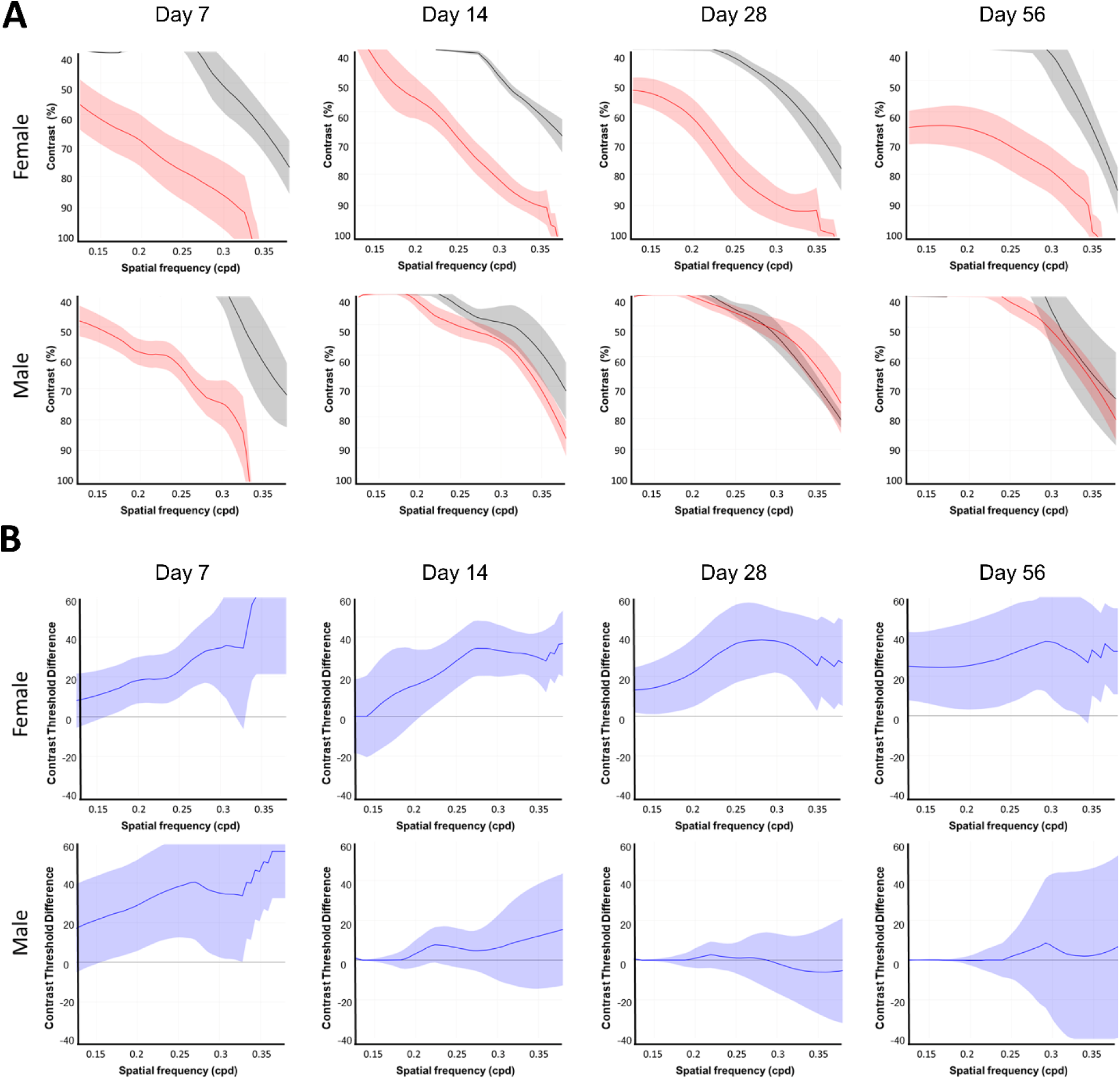
Contrast sensitivity threshold comparisons **A.** Bias-adjusted contrast sensitivity thresholds in sham (black) and TBI (red) groups per time-point in male and female mice based on Fig. 5. Shaded intervals indicate standard error of the mean. **B.** Contrast threshold differences (solid blue line) between sham and TBI mice from (A) with corresponding 95% confidence intervals (shaded blue). In both male and female mice there is a significant decrease in contrast thresholds on day 7 after TBI, which is followed by recovery in a sexually dimorphic manner. In male mice thresholds have recovered by 14 days, whereas in female mice persistent differences are observed up to 56 days post-injury.

Although pVEPs primarily capture cortical processing and do not directly measure SC connectivity, these functional deficits align with the course of anatomical findings of collateral sprouting presented in the previous sections. Together, these results suggest that IA-TBI disrupts the integrity of retinal efferents in a similar manner in male and female mice, but with sexually dimorphic recovery patterns.

### 4. Collateral sprouting does not appear to depend on the Wallerian degeneration of disconnected axons

The mechanisms that drive or facilitate collateral sprouting remain incompletely understood. Nevertheless, Wallerian degeneration of disconnected axons, a process mediated by the NADase SARM1, has been implicated in promoting regeneration within the PNS (Brown et al., 1992, 1994) and heterotopic collateral spouting in the CNS (Steward, 1992; Shi and Stanfield, 1996). In previous studies we demonstrated that, in our TBI model, most RGC axons undergo either immediate or delayed disconnection within the first 24 hours post-injury (Alexandris et al., 2023). Although this disconnection is independent of SARM1 activity, the deletion of *Sarm1* preserves axonal integrity, suppresses microglial reactivity, and delays degeneration for up to three weeks (Alexandris et al., 2023). To investigate whether WD of injured or disconnected axons is required for homotopic collateral sprouting, we assessed changes in CTB labelled retinocollicular terminals of *Sarm1* KO mice after IA-TBI or sham injury. Our findings indicate that *Sarm1* deletion does not significantly alter the extent or progression of collateral sprouting, which was comparable to that observed in wild-type (WT) mice (**Fig. 11; Table S9**).

**Fig. 11.**
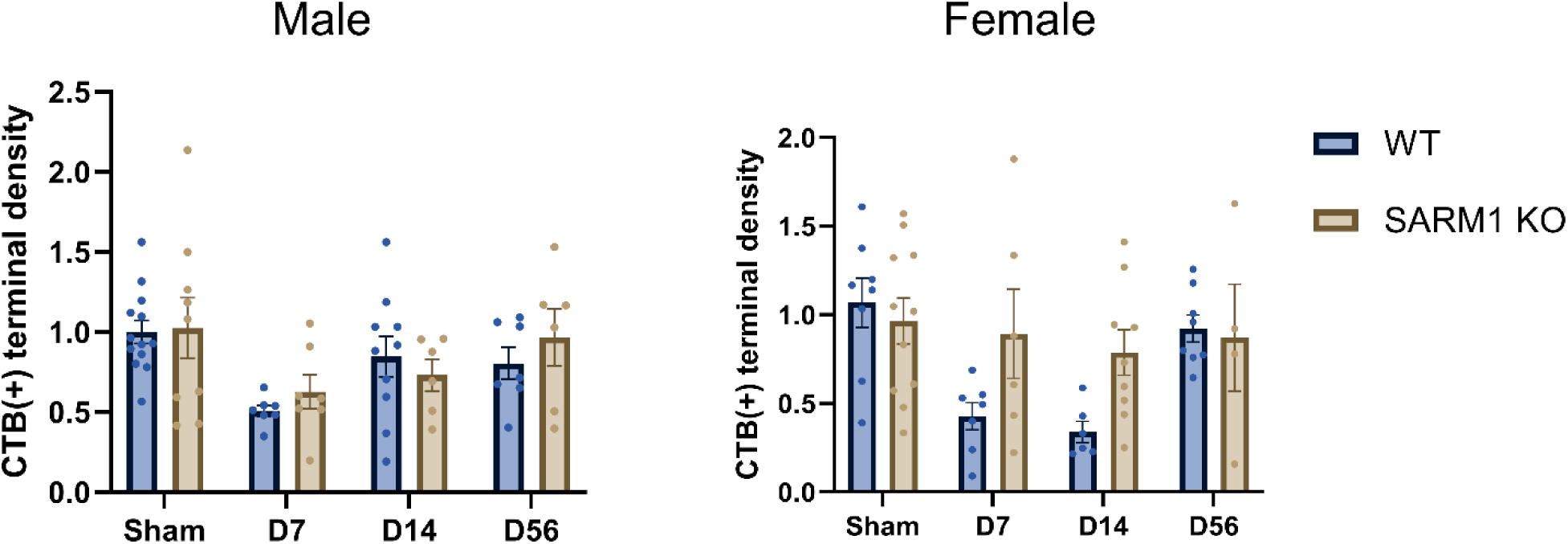
Evidence that Wallerian degeneration is not required for homotopic collateral sprouting. Comparison of CTB(+) axon terminal densities in the superior colliculus of male and female, wt and *Sarm1* KO mice. There are no significant differences between wt and SARM1 KO mice. For details see **supplemental statistical analyses** Table S9.

## Discussion

Our study shows that the adult visual system exhibits a remarkable capacity for structural and functional recovery after traumatic axonal injury. Despite substantial loss of RGC axons and terminals, surviving RGCs achieve collateral sprouting that restores terminal densities to pre-injury levels, with a two-fold increase in terminal arborization per axon. As shown by genetic labeling, terminal axon sprouting is associated with a parallel recovery of synaptic puncta. Structural recovery is accompanied by reestablishment of connectivity as evidenced by transsynaptic tracing and c-Fos mapping, and is associated with an improvement of vision as measured with pVEPs.

Notably, recovery is sexually dimorphic, with female mice demonstrating delayed/ incomplete trajectories of repair and functional recovery. Our work is consistent with a large body of evidence showing robust collateral sprouting responses in the CNS after denervating lesions, but it also uncovers three additional, understudied aspects of CNS repair: the capacity for homotypic sprouting after diffuse lesions in a highly topographically organised circuit (Oswald Steward and Steward, 1991); the reestablishment of connectivity; and the importance of sex as a variable in plasticity after injury.

In contrast to focal or complete lesions, diffuse injuries spare neurons and allow them to process environmental cues needed for rewiring. While homotypic collateral sprouting has been observed before in systems with little topographical organization, such as the ascending dopaminergic pathways (Onn et al., 1986; Blanchard et al., 1996; Finkelstein et al., 2000; Arkadir et al., 2014), we show that homotypic sprouting can occur in a topographically organized circuit and restore connectivity to pre-injury levels. Our observation that surviving RGC axons double their terminal arborization after a ∼50% loss of neighbouring axons is consistent with the principle of proportionality stipulating that the magnitude of sprouting response is commensurate with the degree of deafferentation (Cotman et al., 1981; Steward, 1989)suggesting an intrinsic capacity of a pathway to maintain innervation density based on some homeostatic mechanism (Cotman et al., 1981; Steward, 1989). Although we did not assess the retinotopic specificity of the regenerated connections, our data demonstrate that the gross organization of retinal projections, i.e. laterality, is preserved. Therefore, some basic level of spatial specificity is maintained, with the precise targeting or “fidelity” of these new connections awaiting further exploration.

Functionality of collateral sprouting is an important aspect of recovery and was addressed in this study with complementary approaches. Although c-Fos mapping reveals that stimulated network activity is re-established in the same timeframe as anatomical connectivity, our data indicate that functional recovery is not uniform. The top 10% of neurons show a slower return to baseline activity levels; this delay is especially evident in female mice that do not fully recover even at two months post-injury. This suggests that, while structural aspects of connectivity are promptly restored, some aspects of network function may take longer. The pVEP findings also warrant discussion. Although flash VEPs remain intact (unpublished observations), likely due to redundancy in post-receptor circuits, our pVEP protocol, with its focus on contrast sensitivity, uncovers subtle yet significant functional deficits in traumatic optic axonopathy. In sham animals, pVEP measures show a time-dependent increase in amplitudes, indicative of learning or potentiation. On the other hand, in the IA-TBI groups, regardless of how thresholds are defined, the contrast sensitivity function is compromised, requiring higher contrast levels to evoke cortical responses. Strikingly, these functional deficits differ by sex: while male mice exhibit a recovery to sham-like thresholds by day 14, female mice show persistent deficits, with partial improvement by day 56. These observations suggest that, although repair via sprouting restores a basic anatomical landscape for connectivity—evidenced by both axon tracing and c-Fos mapping— functional recovery, is more complex and may involve additional mechanisms such as activity-dependent synaptic refinement (Murphy and Corbett, 2009). It should be noted here that pVEPs rely on cortical responses largely driven by LGN inputs not directly assessed in our anatomical studies. Therefore, pVEPs serve as a functional proxy of the visual system and not a direct readout of the retinocollicular projection. Regardless, our findings suggest a dissociation between an early structural repair and the late, perhaps more nuanced and sexually dimorphic, recovery of function.

The discovery of a clear sexual dimorphism in recovery trajectories deserves special commentary. Although the acute phase of injury and the immediate degeneration of terminals were similar in both sexes, indicating comparable initial injury burden within the retinofugal pathway, subsequent anatomical and functional recovery diverged markedly. Nonetheless, because we did not assess non-visual systems, sex- or size-dependent differences in injury severity elsewhere cannot be excluded as indirect contributors. Yet, sex-based differences in neuropathological and behavioural outcomes in pre-clinical models of TBI are well recognised (Gupte et al., 2019; Rubin and Lipton, 2019), but the direction of this dimorphism is not straightforward and can vary based on model and outcome. While many studies report a neuroprotective advantage in females with respect to lesion volume, mortality, and acute functional outcomes (Gupte et al., 2019; Rubin and Lipton, 2019), our findings suggest that, when it comes to long-term recovery, females may fare worse. Indeed, persistent deficits in female, but not male, rats have been observed in measures of long-term potentiation in the dentate gyrus (White et al., 2017) and in spatial learning tasks (Wirth et al., 2017) after closed-head injury. These preclinical observations are consistent with clinical evidence indicating that women may experience more severe and enduring symptoms following concussion (Preiss-Farzanegan et al., 2009; Bazarian et al., 2010; Berz et al., 2013; Covassin et al., 2018).

Our findings help clarify patterns of structural axonal and functional recovery and suggest that sex-based differences in recovery trajectories may diverge from those observed typically in the acute phase, pointing to distinct mechanisms underlying long-term outcomes.

Beyond providing valuable insights into plasticity and recovery post TBI, our model serves as a platform for dissecting the relationship between structural and functional repair, and the molecular mechanisms driving collateral sprouting. While WD has been proposed to create a permissive environment for axonal regeneration in the PNS (Brown et al., 1992, 1994) and in heterotypic collateral sprouting in the CNS (Steward, 1992; Shi and Stanfield, 1996), our findings suggest that its role in homotypic collateral sprouting may be limited. This distinction underscores the importance of pathway-specific mechanisms in shaping plasticity. At least based on the hierarchical substitution principle, it is possible that in heterotypic sprouting newly sprouting axons compete with disconnected but preserved axons, whereas in homotypic sprouting this competition may be reduced or absent. Mechanisms other than WD may be more fundamental for enabling homotypic sprouting. Classical studies in entorhinal cortex and the corticospinal tract have demonstrated that collateral sprouting is regulated by neurotrophic factors, cell adhesion and extracellular matrix components (Collazos-Castro and Nieto-Sampedro, 2001; Nicolai Ε. Savaskan and Robert Nitsch, 2001; Hagg, 2006; Thomas Deller et al., 2006). These extrinsic cues define permissive and instructive environments for axonal outgrowth. However, emerging evidence suggests that collateral sprouting may also be driven by intrinsic, context-dependent transcriptional programs distinct from those regulating classical axon regeneration (Li et al., 2010; Carmichael et al., 2017; Lemaitre et al., 2020). Taken together, these findings suggest that collateral sprouting is not simply a passive response to injury-induced vacancy, but may instead require active engagement of specific molecular pathways.

Despite important advances with respect to CNS repair post TBI, our study has some limitations that merit consideration. First, evidence for collateral sprouting, although strong and consistent with previous literature, is primarily inferential. While we demonstrate loss of ∼50% of optic nerve axons ruling out regeneration, we did not visualize branch formation. Nevertheless, collateral sprouting is the only conceivable explanation for the observed restoration of terminals, as evidenced by more than 50% increase in the terminal-to-axon ratio compared to sham-injured mice. In addition, our analysis was performed across the entire RGC population without detailed assessment of microarchitecture of individual RGC terminal fields, leaving open questions about the spatial precision and limits of sprouting. Second, while overall patterns in the LGN match those in the SC, outcomes were not as clear. Our study was principally powered to detect effects in the SC, which may have limited our sensitivity to capture changes in the LGN particularly given the variance and the apparently less robust injury-induced deficits. Differential responses of target neurons to partial denervation may also be possible, weighing against the notion of a generic collateral repair across terminal fields. Third, our study revealed substantial inter-individual variance in retinocollicular terminal density. Although technical factors such as variability in tracer uptake and transport might contribute, nested variance analysis suggests that biological factors predominantly underlie this variance. Previous studies indicate that retinofugal projections and their terminals are subject to activity-dependent remodeling and synaptic refinement both during development and in response to experience (Hooks and Chen, 2006, 2008). Such dynamics may account for individual differences in projection density in the context of circuit plasticity following injury. Recognition of this biological variability is important for interpreting both baseline measures and responses to injury. Fourth, while pVEP recordings provide valuable insights into the integrity of the visual pathway, they do not directly capture the function of subcortical retinofugal circuits.

Unfortunately, behavioral measures such as optomotor reflex responses, which more closely relate to subcortical connectivity, were not proven to be sufficiently sensitive to detect deficits in our hands.

## Conclusion

In conclusion, our study demonstrates that the adult visual system possesses a remarkable capacity for structural and functional recovery after traumatic axonal injury. Surviving RGCs issue collateral sprouting from terminal axons that restores retinocollicular connectivity and at least partially recovers visual function, with notable sex-based differences in recovery trajectories.

These findings illustrate a substantial potential of endogenous repair in the CNS after TBI and underscore the need to further explore the molecular, cellular, and systems-level mechanisms that shape recovery. By establishing a framework to dissect these processes, our work provides a foundation for future studies aimed at enhancing repair strategies in diffuse TBI.

## Supporting information

Supplemental Statistical analyses

## Acknowledgments

We are grateful to Dr. Alex Kolodkin, Professor of Neuroscience (Johns Hopkins University), and Dr. Constantine Frangakis, Professor of Biostatistics (Johns Hopkins University), for insightful discussions. We thank Dr. Victoria Neckles for assistance with optomotor response testing. We also acknowledge Ms. Georgia L. Lawlor and Mr. Sadid Khan for their help in setting up pattern visual evoked potential recordings.

## References

Abbott CJ, Choe TE, Lusardi TA, Burgoyne CF, Wang L, Fortune B (2013) Imaging axonal transport in the rat visual pathway. Biomed Opt Express 4:364–386.

Alexandris AS, Lee Y, Lehar M, Alam Z, McKenney J, Perdomo D, Ryu J, Welsbie D, Zack DJ, Koliatsos VE (2023) Traumatic Axonal Injury in the Optic Nerve: The Selective Role of SARM1 in the Evolution of Distal Axonopathy. J Neurotrauma 40:1743–1761.

Alexandris AS, Lee Y, Lehar M, Alam Z, Samineni P, Tripathi SJ, Ryu J, Koliatsos VE (2022) Traumatic axonopathy in spinal tracts after impact acceleration head injury: Ultrastructural observations and evidence of SARM1-dependent axonal degeneration. Exp Neurol:114252.

Anisimova M, Lamothe-Molina PJ, Franzelin A, Aberra AS, Hoppa MB, Gee CE, Oertner TG (2023) Neuronal FOS reports synchronized activity of presynaptic neurons. :2023.09.04.556168 Available at: https://www.biorxiv.org/content/10.1101/2023.09.04.556168v1 [Accessed March 25, 2025].

Arkadir D, Bergman H, Fahn S (2014) Redundant dopaminergic activity may enable compensatory axonal sprouting in Parkinson disease. Neurology 82:1093–1098.

Bazarian JJ, Blyth B, Mookerjee S, He H, McDermott MP (2010) Sex Differences in Outcome after Mild Traumatic Brain Injury. J Neurotrauma 27:527–539.

Berz K, Divine J, Foss KB, Heyl R, Ford KR, Myer GD (2013) Sex-specific differences in the severity of symptoms and recovery rate following sports-related concussion in young athletes. Phys Sportsmed 41:58–63.

Blanchard V, Anglade P, Dziewczapolski G, Savasta M, Agid Y, Raisman-Vozari R (1996) Dopaminergic sprouting in the rat striatum after partial lesion of the substantia nigra. Brain Res 709:319–325.

Brown MC, Lunn ER, Perry VH (1992) Consequences of slow Wallerian degeneration for regenerating motor and sensory axons. J Neurobiol 23:521–536.

Brown MC, Perry VH, Hunt SP, Lapper SR (1994) Further studies on motor and sensory nerve regeneration in mice with delayed Wallerian degeneration. Eur J Neurosci 6:420–428.

Buzsaki G, Mizuseki K (2014) The log-dynamic brain: how skewed distributions affect network operations. Nat Rev Neurosci 15:264–278.

Byun H, Kwon S, Ahn H-J, Liu H, Forrest D, Demb JB, Kim I-J (2016) Molecular features distinguish ten neuronal types in the mouse superficial superior colliculus. J Comp Neurol 524:2300–2321.

Carmichael ST, Kathirvelu B, Schweppe CA, Nie EH (2017) Molecular, cellular and functional events in axonal sprouting after stroke. Exp Neurol 287:384–394.

Chaudhuri A, Zangenehpour S, Rahbar-Dehgan F, Ye F (2000) Molecular maps of neural activity and quiescence. Acta Neurobiol Exp (Wars) 60:403–410.

Collazos-Castro JE, Nieto-Sampedro M (2001) Developmental and reactive growth of dentate gyrus afferents: Cellular and molecular interactions. Restorative Neurology and Neuroscience 19:169–187.

Cotman CW, Nieto-Sampedro M, Harris EW (1981) Synapse replacement in the nervous system of adult vertebrates. Physiological Reviews 61:684–784.

Covassin T, Savage JL, Bretzin AC, Fox ME (2018) Sex differences in sport-related concussion long-term outcomes. Int J Psychophysiol 132:9–13.

Efron B, Stein C (1981) The Jackknife Estimate of Variance. The Annals of Statistics 9:586–596.

Ellis EM, Gauvain G, Sivyer B, Murphy GJ (2016) Shared and distinct retinal input to the mouse superior colliculus and dorsal lateral geniculate nucleus. J Neurophysiol 116:602–610.

Exner S (1885) Notiz zu der Frage von der Faservertheilung mehrerer Nerven in einem Muskel. Pflüger, Arch 36:572–576.

Fawcett JW (2020) The Struggle to Make CNS Axons Regenerate: Why Has It Been so Difficult? Neurochem Res 45:144–158.

Finkelstein DI, Stanic D, Parish CL, Tomas D, Dickson K, Horne MK (2000) Axonal sprouting following lesions of the rat substantia nigra. Neuroscience 97:99–112.

Guo YP, Sun X, Li C, Wang NQ, Chan Y-S, He J (2007) Corticothalamic synchronization leads to c-fos expression in the auditory thalamus. Proc Natl Acad Sci U S A 104:11802–11807.

Gupte R, Brooks W, Vukas R, Pierce J, Harris J (2019) Sex Differences in Traumatic Brain Injury: What We Know and What We Should Know. J Neurotrauma 36:3063–3091.

Hagg T (2006) Collateral Sprouting as a Target for Improved Function after Spinal Cord Injury. Journal of Neurotrauma 23:281–294.

Hooks BM, Chen C (2006) Distinct roles for spontaneous and visual activity in remodeling of the retinogeniculate synapse. Neuron 52:281–291.

Hooks BM, Chen C (2008) Vision triggers an experience-dependent sensitive period at the retinogeniculate synapse. J Neurosci 28:4807–4817.

Lemaitre D, Hurtado ML, De Gregorio C, Oñate M, Martínez G, Catenaccio A, Wishart TM, Court FA (2020) Collateral Sprouting of Peripheral Sensory Neurons Exhibits a Unique Transcriptomic Profile. Mol Neurobiol 57:4232–4249.

Li S, Overman JJ, Katsman D, Kozlov SV, Donnelly CJ, Twiss JL, Giger RJ, Coppola G, Geschwind DH, Carmichael ST (2010) An age-related sprouting transcriptome provides molecular control of axonal sprouting after stroke. Nat Neurosci 13:1496–1504.

Liu C, Alexandris AS, Gharagozloo M, Garton T, Calabresi PA, Koliatsos VE (2025) Temporal progression of axonal degeneration in the Visual System in EAE: Insights from high-resolution neuropathology. JNEN.

Makin TR, Krakauer JW (2023) Against cortical reorganisation Pruszynski JA, de Lange FP, eds. eLife 12:e84716.

Martersteck EM, Hirokawa KE, Evarts M, Bernard A, Duan X, Li Y, Ng L, Oh SW, Ouellette B, Royall JJ, Stoecklin M, Wang Q, Zeng H, Sanes JR, Harris JA (2017) Diverse Central Projection Patterns of Retinal Ganglion Cells. Cell Reports 18:2058–2072.

Nicolai Ε. Savaskan, Robert Nitsch (2001) Molecules Involved in Reactive Sprouting in the Hippocampus. Reviews in the Neurosciences 12:195–216.

Onn SP, Berger TW, Stricker EM, Zigmond MJ (1986) Effects of intraventricular 6-hydroxydopamine on the dopaminergic innervation of striatum: histochemical and neurochemical analysis. Brain Res 376:8–19.

Oswald Steward, Steward O (1991) Synapse Replacement on Cortical Neurons following Denervation. :81–131.

Povlishock JT (1992) Traumatically induced axonal injury: pathogenesis and pathobiological implications. Brain Pathol 2:1–12.

Preiss-Farzanegan SJ, Chapman B, Wong TM, Wu J, Bazarian JJ (2009) The relationship between gender and postconcussion symptoms after sport-related mild traumatic brain injury. PM R 1:245–253.

Rubin TG, Lipton ML (2019) Sex Differences in Animal Models of Traumatic Brain Injury. J Exp Neurosci 13:1179069519844020.

Schindelin J, Arganda-Carreras I, Frise E, Kaynig V, Longair M, Pietzsch T, Preibisch S, Rueden C, Saalfeld S, Schmid B, Tinevez JY, White DJ, Hartenstein V, Eliceiri K, Tomancak P, Cardona A (2012) Fiji: an open-source platform for biological-image analysis. Nat Methods 9:676–682.

Shi B, Stanfield BB (1996) Differential sprouting responses in axonal fiber systems in the dentate gyrus following lesions of the perforant path in WLDs mutant mice. Brain Res 740:89–101.

Steward O (1989) Reorganization of neuronal connections following CNS trauma: principles and experimental paradigms. J Neurotrauma 6:99–152.

Steward O (1992) Signals that induce sprouting in the central nervous system: sprouting is delayed in a strain of mouse exhibiting delayed axonal degeneration. Exp Neurol 118:340–351.

Stoica BA, Faden AI (2010) Cell death mechanisms and modulation in traumatic brain injury. Neurotherapeutics 7:3–12.

Sulaiman W, Gordon T (2013) Neurobiology of Peripheral Nerve Injury, Regeneration, and Functional Recovery: From Bench Top Research to Bedside Application. Ochsner J 13:100–108.

Thomas Deller, Deller T, Carola A. Haas, Haas CA, Thomas M. Freiman, Thomas M. Freiman, Freiman TM, Amie L. Phinney, Phinney AL, Mathias Jucker, Jucker M, Michael Frotscher, Frotscher M (2006) Lesion-induced axonal sprouting in the central nervous system. Advances in Experimental Medicine and Biology 557:101–121.

Tsai NY, Wang F, Toma K, Yin C, Takatoh J, Pai EL, Wu K, Matcham AC, Yin L, Dang EJ, Marciano DK, Rubenstein JL, Wang F, Ullian EM, Duan X (2022) Trans-Seq maps a selective mammalian retinotectal synapse instructed by Nephronectin. Nat Neurosci 25:659–674.

Uccellini MB, Bardina SV, Sánchez-Aparicio MT, White KM, Hou YJ, Lim JK, García-Sastre A (2020) Passenger Mutations Confound Phenotypes of SARM1-Deficient Mice. Cell Rep 31:107498.

White ER, Pinar C, Bostrom CA, Meconi A, Christie BR (2017) Mild Traumatic Brain Injury Produces Long-Lasting Deficits in Synaptic Plasticity in the Female Juvenile Hippocampus. J Neurotrauma 34:1111–1123.

Wirth P, Yu W, Kimball AL, Liao J, Berkner P, Glenn MJ (2017) New method to induce mild traumatic brain injury in rodents produces differential outcomes in female and male Sprague Dawley rats. J Neurosci Methods 290:133–144.

Wood SN (2011) Fast Stable Restricted Maximum Likelihood and Marginal Likelihood Estimation of Semiparametric Generalized Linear Models. Journal of the Royal Statistical Society Series B: Statistical Methodology 73:3–36.

Xu L, Nguyen JV, Lehar M, Menon A, Rha E, Arena J, Ryu J, Marsh-Armstrong N, Marmarou CR, Koliatsos VE (2016) Repetitive mild traumatic brain injury with impact acceleration in the mouse: Multifocal axonopathy, neuroinflammation, and neurodegeneration in the visual system. Exp Neurol 275 Pt 3:436–449.

Yassin L, Benedetti BL, Jouhanneau J-S, Wen JA, Poulet JFA, Barth AL (2010) An embedded subnetwork of highly active neurons in the neocortex. Neuron 68:1043–1050.

Ziogas NK, Koliatsos VE (2018) Primary Traumatic Axonopathy in Mice Subjected to Impact Acceleration: A Reappraisal of Pathology and Mechanisms with High-Resolution Anatomical Methods. J Neurosci 38:4031–4047.

