## Supplemental Figures for "Recovery of retinal terminal fields after traumatic brain injury: evidence of collateral sprouting and sexual dimorphism"

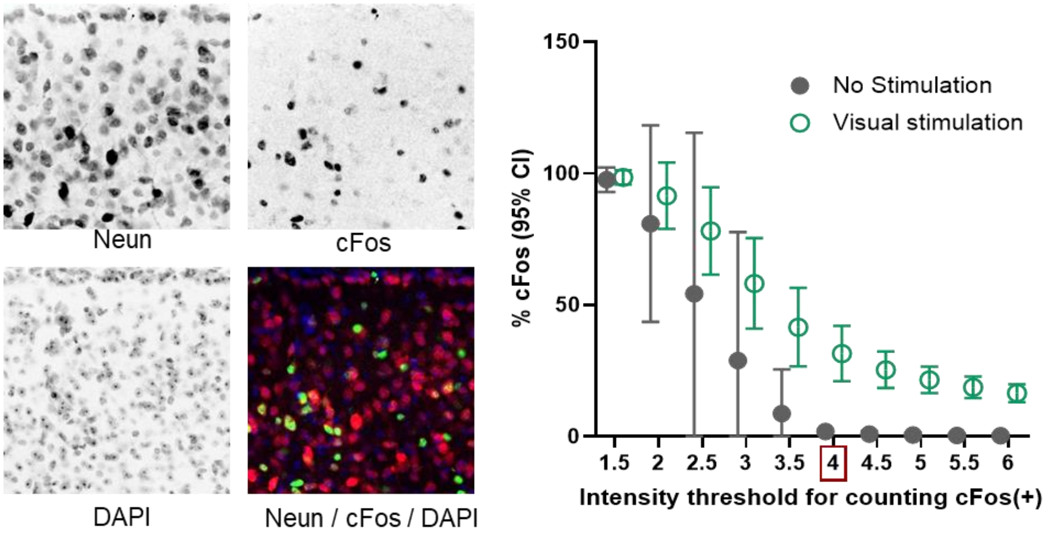


**Fig. S1**. Determination of c-Fos intensity threshold for connectivity analysis. **A.** Confocal image through the superior colliculus showing immunoreactivity for NeuN and c-Fos. **B.** Reverse cumulative frequency distribution showing fraction (%) of c-Fos(+)/NeuN(+) cells for each intensity threshold in visually stimulated (green) and light deprived mice (grey). The minimum threshold which separates light deprived from visually stimulated conditions is highlighted (red rectangle).

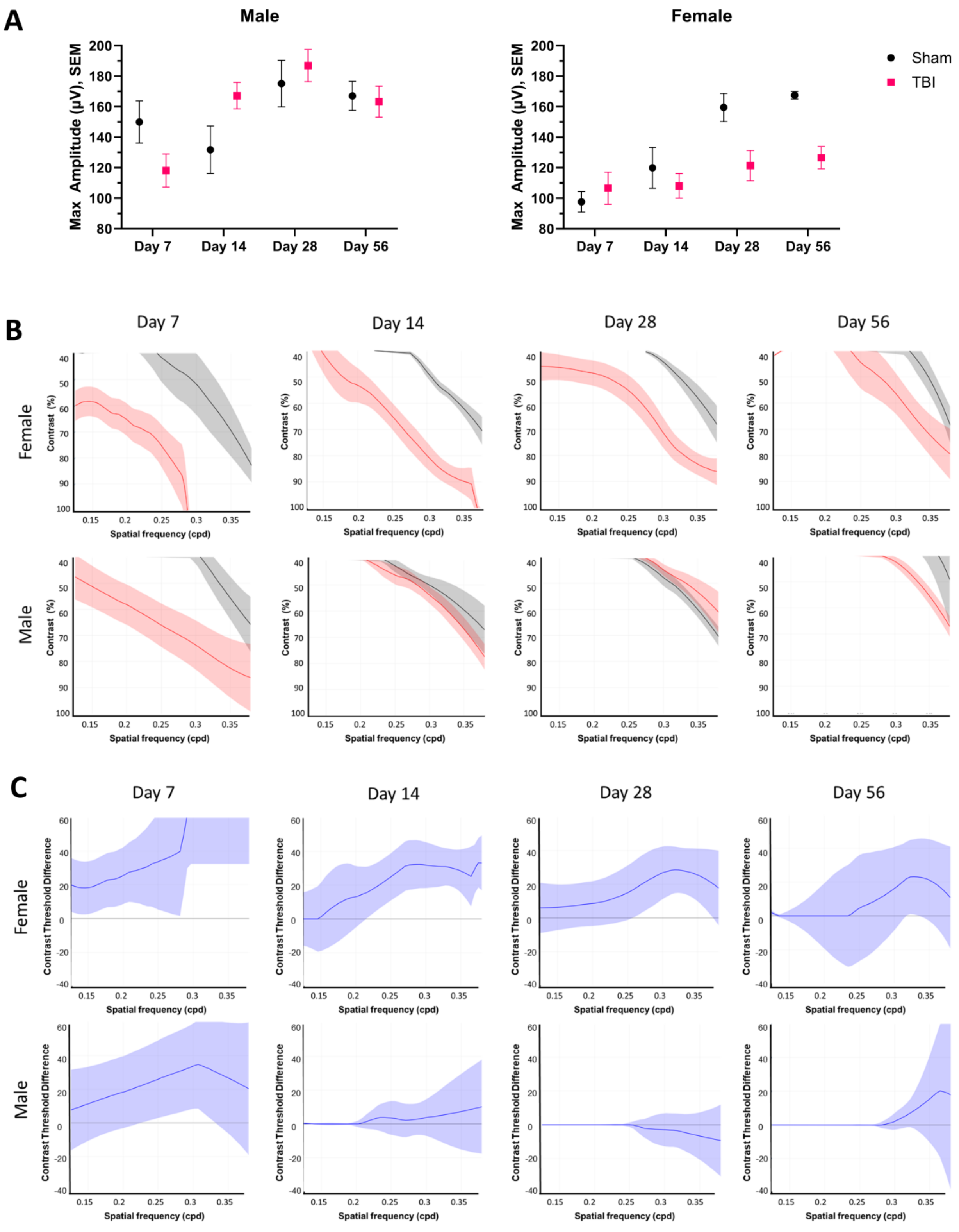

A
Male
Female
Max Amplitude (μV), SEM
Day 7
Day 14
Day 28
Day 56
Sham
TBI
B
Female
Male
Contrast (%)
Spatial frequency (cpd)
C
Contrast Threshold Difference

**Fig. S2.** Contrast sensitivity threshold comparisons based on a global fixed threshold (defined as 50% of maximum amplitude for sham male and female mice at day 7) **A**. Bias-adjusted max amplitude averages per group were derived from the heatmaps in Figure 8D-E and plotted across time. See **Supplemental Statistical Analyses** Table S7 for details. **B.** Bias-adjusted contrast sensitivity thresholds in sham (black) and TBI (red) groups per time-point in male and female mice. Shaded intervals indicate standard error of the mean. **C.** Contrast threshold differences (solid blue line) between sham and TBI mice from (A) with corresponding 95% confidence intervals (shaded blue).
