## Supplemental Statistical analyses for "Recovery of retinal terminal fields after traumatic brain injury: evidence of collateral sprouting and sexual dimorphism"

**Table S1.** Statistical analyses related to Fig. 1: Course of collateral sprouting in retinocollicular projections after IA-TBI in male and female mice.

| **Fig 1D.** Changes in CTB(+) terminal density after IA-TBI | | |
|---|---|---|
| **Two-way ANOVA** | **Post-hoc comparisons** | |
| *Sex*<br>*F(1, 69) = 2.154; p=0.1468*<br>*Injury groups*<br>*F(4, 69) = 10.76; p<0.0001*<br><br>*Interaction*<br>*F(4, 69) = 3.062; p=0.0221* | Male | |
| | D7 vs. Sham | **t(69)=3.788; p=0.0013** |
| | D7 vs. D14 | **t(69)=2.48; p=0.046** |
| | D7 vs. D28 | **t(69)=2.318; p=0.0463** |
| | D7 vs. D56 | **t(69)=1.998; p=0.0496** |
| | Female | |
| | D7 vs. Sham | **t(69)=4.465; p=0.0001** |
| | D7 vs. D14 | t(69)=0.5736; p=0.5681 |
| | D7 vs. D28 | **t(69)=2.499; p=0.0295** |
| | D7 vs. D56 | **t(69)=3.45; p=0.0029** |

| **Fig 1E.** Changes in RGC densities and the CTB/RGC ratio (pooled sex analysis) | | |
|---|---|---|
| **One-way ANOVA** | **Post-hoc comparisons** | |
| **RGC Densities** | | |
| F (2, 39) = 10.94; p=0.0002 | Sham vs. Day 7 | **t(39)=2.81; p=0.0153** |
| | Sham vs. Day 56 | **t(39)=4.643; p=0.0001** |
| | Day 7 vs. Day 56 | **t(39)=1.833; p=0.0745** |
| **CTB/RGC Ratio** | | |
| F (2, 39) = 19.56; P<0.0001 | Sham vs. Day 7 | **t(39)=6.634; p<0.0001** |
| | Sham vs. Day 56 | t(39)=1.751; p=0.4384 |
| | Day 7 vs. Day 56 | **t(39)=8.385; p<0.0001** |

| **Fig 1F.** Changes in Axon counts and the CTB/Axon count ratio (pooled sex analysis) | | |
|---|---|---|
| **One-way ANOVA** | **Post-hoc comparisons** | |
| **Axon counts** | | |
| F (2, 10) = 13.01; p=0.0017 | Sham vs. Day 7 | t(10)=0.3903; p=0.7045 |
| | Sham vs. Day 56 | **t(10)=4.729; p=0.0024** |
| | Day 7 vs. Day 56 | **t(10)=4.116; p=0.0042** |
| **CTB/Axon Ratio** | | |
| F (2, 9) = 14.14; p=0.0017 | Sham vs. Day 7 | **t(9)=2.58; p=0.0454** |
| | Sham vs. Day 56 | **t(9)=2.737; p=0.0454** |
| | Day 7 vs. Day 56 | **t(9)=5.317; p=0.0014** |

**Table S2.** Statistical analyses related to Fig. 4: Course of collateral sprouting in retinocollicular projections after IA-TBI using genetic tracers.

| **Fig. 3D.** Changes in tdTomato(+) terminal axon densities, SypGFP(+) synaptic densities and ratios of synapses per terminal axon arbors. | | |
|---|---|---|
| **One-way ANOVA** | **Post-hoc comparisons** | |
| **Terminal axon density** | | |
| F (2, 11) = 4.916 p=0.0298 | Day 7 vs. Sham | **t(11)=2.274; p=0.044** |
| | Day 7 vs. Day 28 | **t(11)=2.976; p=0.025** |
| **SypGFP density** | | |
| F (2, 11) = 9.671; p=0.0038 | Day 7 vs. Sham | **t(11)=3.78; p=0.0061** |
| | Day 7 vs. Day 28 | **t(11)=3.764; p=0.0061** |
| **SypGFP per terminal arbor** | | |
| F (2, 11) = 2.626; p=0.1168 | Day 7 vs. Sham | t(11)=0.999; p=0.3393 |
| | Day 7 vs. Day 28 | **t(11)=2.29; p=0.0837** |

**Table S3.** Statistical analyses related to Fig. 5: Transsynaptic-tracing based analysis of retinocollicular connectivity after IA-TBI.

| **Fig. 5D.** Changes in retinocollicular connectivity based on WGA-mCherry labeling of Neun(+) SC neurons in male and female mice. | | |
|---|---|---|
| **Two-way ANOVA** | **Post-hoc comparisons (main effect – pooled sex analysis)** | |
| Sex<br>*F (1, 39) = 0.6989; p=0.4083*<br>Injury groups<br>*F (3, 39) = 5.306; p=0.0036*<br><br>Interaction<br>*F (1, 39) = 0.6989; p=0.4083* | Sham vs. Day 7 | **t(39)=2.977; p=0.0246** |
| | Sham vs. Day 14 | t(39)=2.477; p=0.0521 |
| | Sham vs. Day 56 | t(39)=0.1146; p=0.9094 |
| | Day 7 vs. Day 14 | t(39)=0.8391; p=0.6478 |
| | Day 7 vs. Day 56 | **t(39)=3.127; p=0.0198** |
| | Day 14 vs. Day 56 | **t(39)=2.646; p=0.046** |

**Table S4.** Statistical analyses related to Fig. 6: cFos based analysis of retinocollicular connectivity after IA-TBI.

**Fig 6. C.** Changes in retinocollicular connectivity based on cFos expression in Neun(+) SC neurons in male and female mice.

| Two-way ANOVA | Post-hoc comparisons | |
|---|---|---|
| | **Males** | |
| | Sham vs D56 | t(16)=0.5546; p=0.5868 |
| Sex | Sham vs. D7 | **t(16)=3.449; p=0.0131** |
| *F (1, 33) = 0.6955; p=0.5614* | D7 vs. D14 | t(16)=2.071; p=0.1067 |
| Injury groups | D7 vs. D56 | **t(16)=2.894; p=0.0314** |
| *F (3, 33) = 17.97; p<0.0001* | **Females** | |
| Interaction | **Sham vs D56** | **t(17)=3.134; p=0.0121** |
| *F (3, 33) = 0.6955; p=0.5614* | **Sham vs. D7** | **t(17)=8.001; p=<0.0001** |
| | **D7 vs. D14** | **t(17)=2.691; p=0.0154** |
| | **D7 vs. D56** | **t(17)=4.506; p=0.0009** |

**Fig 6. E** Changes in median cFos intensities in the top 10% of neurons.

| Two-way ANOVA | Post-hoc comparisons | |
|---|---|---|
| | **Males** | |
| | NS vs. Sham | **t(17)=2.896; p=0.0335** |
| Sex | Sham vs. D7 | **t(17)=2.317; p=0.0335** |
| *F (1, 33) = 0.04474; p=0.8338* | D7 vs. D14 | **t(17)=2.976; p=0.0335** |
| Injury groups | D7 vs. D56 | **t(17)=2.931; p=0.0335** |
| *F (3, 33) = 9.692; p<0.0001* | **Females** | |
| Interaction | NS vs. Sham | **t(18)=4.763; p=0.0005** |
| *F (3, 33) = 2.648; p=0.0652* | Sham vs. D7 | **t(18)=5.275; p=0.0002** |
| | D7 vs. D14 | **t(18)=1.883; p=0.076** |
| | D7 vs. D56 | **t(18)=2.715; p=0.0282** |

**Table S5.** Statistical analyses related to Fig. 7: Analyses of retinogeniculate sprouting and connectivity

| **Figure 7. A**. CTB(+) terminal densities of the contralateral dorsal and ventral lateral geniculate nucleus (dLG and vLG) | | |
|---|---|---|
| **Two-way ANOVA** | **Post-hoc comparisons (main effect – pooled sex analysis))** | |
| **dLGN** | | |
| Sex<br>*F (1, 29) = 1.290 p=0.2654*<br>Injury groups<br>*F (2, 29) = 2.499 p=0.0997*<br><br>Interaction<br>*F (2, 29) = 0.9442 p=0.4006* | Sham vs. Day 7 | t(29)=2.234; p=0.0968 |
| | Sham vs. Day 56 | t(29)=1.36; p=0.3344 |
| | Day 7 vs. Day 56 | t(29)=0.792; p=0.4348 |
| **vLGN** | | |
| Sex<br>*F (1, 30) = 0.02391 p=0.8781*<br>Injury groups<br>*F (2, 30) = 1.106 p=0.3441*<br><br>Interaction<br>*F (2, 30) = 0.07065 p=0.9319* | Sham vs. Day 7 | t(30)=1.15; p=0.4513 |
| | Sham vs. Day 56 | t(30)=1.437; p=0.4094 |
| | Day 7 vs. Day 56 | t(30)=0.413; p=0.6826 |

| **Figure 7. B**. Transynaptic-tracing based connectivity analysis in the dLG and vLG | | |
|---|---|---|
| **Two-way ANOVA** | **Post-hoc comparisons (main effect – pooled sex analysis)** | |
| **dLGN** | | |
| Sex<br>*F (1, 35) = 0.1640 p=0.6880*<br>Injury groups<br>*F (3, 35) = 8.810 p=0.0002*<br><br>Interaction<br>*F (3, 35) = 2.268 p=0.0977* | **Day 7 vs. Sham** | **t(35)=3.824; p=0.0016** |
| | Day 7 vs. Day 14 | t(35)=0.4271; p=0.6719 |
| | Day 7 vs. Day 56 | t(35)=1.295; p=0.3661 |
| **vLGN** | | |
| Sex<br>*F (1, 39) = 0.5031 p=0.4824*<br>Injury groups<br>*F (3, 39) = 4.411 p=0.0092*<br><br>Interaction<br>*F (3, 39) = 0.6630 p=0.5798* | **Day 7 vs. Sham** | **t(39)=3.107; p=0.0105** |
| | Day 7 vs. Day 14 | t(39)=0.8664; p=0.3916 |
| | **Day 7 vs. Day 56** | **t(39)=2.461; p=0.0365** |

| **Figure 7. C**. c-Fos based connectivity analysis in the dLG and vLG. | | |
|---|---|---|
| **Two-way ANOVA** | **Post-hoc comparisons (main effect – pooled sex analysis)** | |
| **dLGN** | | |
| Sex<br>*F (1, 33) = 1.251 p=0.2714* | Day 7 vs. NS | t(33)=1.632; p=0.2118 |

| | | |
|---|---|---|
| Injury groups<br>*F (4, 33) = 5.062* *p=0.0027*<br><br>Interaction<br>*F (4, 33) = 0.8457* *p=0.5064* | **Day 7 vs. Sham** | **t(33)=3.012; p=0.0197** |
| | Day 7 vs. Day 14 | t(33)=0.453; p=0.6535 |
| | Day 7 vs. Day 56 | t(33)=1.861; p=0.2001 |
| **vLGN** | | |
| Sex<br>*F (1, 33) = 0.03442* *p=0.8539*<br>Injury groups<br>*F (4, 33) = 4.785* *p=0.0037*<br><br>Interaction<br>*F (4, 33) = 2.111* *p=0.1016* | Day 7 vs. NS | t(33)=2.475; p=0.0725 |
| | Day 7 vs. Sham | t(33)=2.253; p=0.0903 |
| | Day 7 vs. Day 14 | t(33)=0.9229; p=0.4335 |
| | Day 7 vs. Day 56 | t(33)=1.178; p=0.4335 |

**Table S6.** Statistical analyses related to Fig.8 B-C.

| **Two-way ANOVA** | **Post-hoc comparisons (TBI vs Sham)** | |
|---|---|---|
| **Male scotopic a-wave** | | |
| Time<br>*F (1.615, 20.99) = 1.465; p=0.2515*<br>Injury groups<br>*F (1, 13) = 0.4868; p=0.4976*<br>Interaction<br>*F (3, 39) = 0. ; p=0.7193* | Baseline | t(12.92)=0.9395; p=(0.7436) |
| | D7 | t(8.225)=0.3553; p=(0.7436) |
| | D14 | t(9.497)=1.397; p=(0.5781) |
| | D28 | t(13)=0.9017; p=(0.7436) |
| **Female scotopic a-wave** | | |
| Time<br>*F (1.668, 21.68) = 18.40; p<0.0001*<br>Injury groups<br>*F (1, 13) = 0.3145; p=0.5845*<br>Interaction<br>*F (3, 39) = 0.1994; p==0.8961* | Baseline | t(10.51)=0.7338; p=(0.9264) |
| | D7 | t(11.43)=0.4867; p=(0.9824) |
| | D14 | t(12.95)=1.01; p=(0.7996) |
| | D28 | t(11.86)=0.7396; p=(0.9234) |
| **Male scotopic b-wave** | | |
| Time<br>*F (2.363, 30.72) = 0.472; p=0.6596*<br>Injury groups<br>*F (1, 13) = 2.124; p=0.1687*<br>Interaction<br>*F (3, 39) = 3.009; p=0.0417* | Baseline | t(10.7)=1.548; p=(0.3874) |
| | D7 | t(12.57)=0.09284; p=(0.9275) |
| | D14 | t(10.02)=2.933; p=(0.0584) |
| | D28 | t(9.689)=1.445; p=(0.3874) |
| **Female scotopic b-wave** | | |
| Time<br>*F (1.668, 21.68) = 18.40; p<0.0001*<br>Injury groups<br>*F (1, 13) = 0.3145; p=0.5845*<br>Interaction<br>*F (3, 39) = 0.1994; p=0.8961* | Baseline | t(11.87)=0.158; p=(0.9623) |
| | D7 | t(9.754)=0.6877; p=(0.9412) |
| | D14 | t(10.01)=0.2524; p=(0.9623) |
| | D28 | t(8.939)=0.4754; p=(0.9556) |
| **Male photopic a-wave** | | |
| Time<br>*F (2.473, 32.15) = 0.7427; p=0.5104*<br>Injury groups<br>*F (1, 13) = 1.906; p=P=0.1907*<br>Interaction<br>*F (3, 39) = 1.314; p=0.2836* | Baseline | t(12.92)=0.9395; p=(0.7436) |
| | D7 | t(8.225)=0.3553; p=(0.7436) |
| | D14 | t(9.497)=1.397; p=(0.5781) |
| | D28 | t(13)=0.9017; p=(0.7436) |
| **Female photopic a-wave** | | |
| Time<br>*F (2.491, 32.38) = 11.05; p<0.0001* | Baseline | t(12.58)=2.705; p=(0.0718) |
| | D7 | t(7.442)=0.4546; p=(0.6623) |

| | | |
|---|---|---|
| Injury groups<br>*F (1, 13) = 2.405; p=0.1450*<br>Interaction<br>*F (3, 39) = 4.436; p=0.0089* | D14 | t(12.2)=2.099; p=(0.1623) |
| | D28 | t(12.48)=1.835; p=(0.1726) |
| **Male photopic b-wave** | | |
| Time<br>*F (2.531, 32.91) = 1.785; p=0.1761*<br>Injury groups<br>*F (1, 13) = 8.585; p=0.0117*<br>Interaction<br>*F (3, 39) = 1.508; p=0.2276* | Baseline | t(10.62)=0.3235; p=(0.8512) |
| | D7 | t(12.06)=2.418; p=(0.1233) |
| | D14 | t(9.851)=0.5205; p=(0.8512) |
| | D28 | t(9.146)=2.465; p=(0.1233) |
| **Female photopic b-wave** | | |
| Time<br>*F (2.194, 28.52) = 2.435; p=0.1012*<br>Injury groups<br>*F (1, 13) = 0.6962; p=.4191*<br>Interaction<br>*F (3, 39) = 0.8375; p=0.4815* | Baseline | t(12.99)=0.7403; p=(0.853) |
| | D7 | t(10.25)=0.1162; p=(0.9862) |
| | D14 | t(12.72)=0.1507; p=(0.9862) |
| | D28 | t(11.38)=1.597; p=(0.4472) |
| **Male 10 Hz Flicker N1-P1 wave** | | |
| Time<br>*F (2.730, 35.49) = 4.498; p=0.0107*<br>Injury groups<br>*F (1, 13) = 1.751; p=0.2086*<br>Interaction<br>*F (3, 39) = 1.162; p==0.3364* | Baseline | t(12.36)=0.5567; p=(0.8234) |
| | D7 | t(11.99)=0.699; p=(0.8234) |
| | D14 | t(7.924)=0.8148; p=(0.8234) |
| | D28 | t(8.852)=2.055; p=(0.2538) |
| **Female 10 Hz Flicker N1-P1 wave** | | |
| Time<br>*F (2.318, 30.13) = 5.456; p=0.0071*<br>Injury groups<br>*F (1, 13) = 1.517; p=0.2398*<br>Interaction<br>*F (3, 39) = 0.03957; p=0.9893* | Baseline | t(12.32)=0.5914; p=(0.8107) |
| | D7 | t(11.92)=1.186; p=(0.6983) |
| | D14 | t(9.095)=1.031; p=(0.6983) |
| | D28 | t(10.49)=0.5878; p=(0.8107) |

**Table S7.** Generative Additive Model parameters corresponding to panels in Fig. 9D-E. GAMs were fitted for each group per time point and sex. $R^2$ indicates the proportion of variance in the amplitudes explained by the model. Effective degrees of freedom (EDF) for fixed effects (parametric), smooth functions (contrast, spatial frequency and their interaction) and residual are shown. Residual EDF was higher in TBI mice compared to sham, suggesting greater variability in individual responses. Despite this, $R^2$ values remained moderate to high (0.5–0.7), indicating that the model captures key trends despite measurement variability.

| **Cohorts** | | | **GAM parameters** | | | | | | |
|---|---|---|---|---|---|---|---|---|---|
| | | | | **Effective degrees of freedom** | | | | | |
| **Sex** | **Group** | **Day** | **$R^2$** | **Parametric** | **Contrast** | **Spatial Frequency** | **Contrast * Spatial Frequency** | **Total** | **Residual** |
| Female | Sham | 7 | 0.40 | 1 | 1.41 | 1.74 | 1.45 | 5.59 | 78.4 |
| | | 14 | 0.60 | 1 | 1.58 | 1.89 | 1 | 5.48 | 50.5 |
| | | 28 | 0.74 | 1 | 1.74 | 1.93 | 1.63 | 6.30 | 49.7 |
| | | 56 | 0.64 | 1 | 1.57 | 1.92 | 1 | 5.49 | 36.5 |
| | TBI | 7 | 0.65 | 1 | 1.82 | 1.36 | 1 | 5.18 | 106.8 |
| | | 14 | 0.56 | 1 | 1 | 1.74 | 1 | 4.74 | 93.3 |
| | | 28 | 0.55 | 1 | 1 | 1.47 | 1.78 | 5.25 | 92.8 |
| | | 56 | 0.59 | 1 | 1 | 1.83 | 1 | 4.83 | 51.2 |
| Male | Sham | 7 | 0.51 | 1 | 1 | 1.66 | 1 | 4.66 | 65.3 |
| | | 14 | 0.52 | 1 | 1.77 | 1.73 | 1 | 5.50 | 64.5 |
| | | 28 | 0.71 | 1 | 1.81 | 1.82 | 1 | 5.64 | 64.4 |
| | | 56 | 0.40 | 1 | 1 | 1.65 | 1 | 4.66 | 65.3 |
| | TBI | 7 | 0.37 | 1 | 1 | 1 | 1 | 4.00 | 108.0 |
| | | 14 | 0.65 | 1 | 1 | 1.91 | 1.72 | 5.63 | 106.4 |
| | | 28 | 0.61 | 1 | 1.89 | 1.51 | 1.77 | 6.17 | 105.8 |
| | | 56 | 0.61 | 1 | 1.8 | 1.96 | 1 | 5.76 | 106.2 |

**Table S8.** Statistical analyses related to Fig. S2.A: Maximum pattern reversal VEP amplitudes change across time.

| Two-way ANOVA | Post-hoc comparisons | |
|---|---|---|
| **Male** | | |
| Time<br>F (3, 44) = 5.828; p=0.0019<br>Injury<br>F (1, 44) = 0.1209; p=0.7297<br>Interaction<br>F (3, 44) = 2.831; p=0.0492 | **Sham** | |
| | Day 7 vs. Day 14 | t(44)=0.9791; p=0.7031 |
| | Day 7 vs. Day 28 | t(44)=1.357; p=0.5518 |
| | Day 7 vs. Day 56 | t(44)=0.9229; p=0.7031 |
| | Day 14 vs. Day 28 | t(44)=2.336; p=0.1363 |
| | Day 14 vs. Day 56 | t(44)=1.902; p=0.2805 |
| | Day 28 vs. Day 56 | t(44)=0.4338; p=0.7031 |
| | **TBI** | |
| | **Day 7 vs. Day 14** | **t(44)=3.335; p=0.0087** |
| | **Day 7 vs. Day 28** | **t(44)=4.677; p=0.0002** |
| | **Day 7 vs. Day 56** | **t(44)=3.069; p=0.0146** |
| | Day 14 vs. Day 28 | t(44)=1.343; p=0.3378 |
| | Day 14 vs. Day 56 | t(44)=0.2654; p=0.7919 |
| | Day 28 vs. Day 56 | t(44)=1.608; p=0.3068 |
| **Female** | | |
| Time<br>F (3, 35) = 8.834; p=0.0002<br>Injury<br>F (1, 35) = 7.711; p=0.0088<br>Interaction<br>F (3, 35) = 2.854; p=0.0511 | **Sham** | |
| | Day 7 vs. Day 14 | t(35)=1.51; p=0.2605 |
| | **Day 7 vs. Day 28** | **t(35)=4.191; p=0.0009** |
| | **Day 7 vs. Day 56** | **t(35)=4.321; p=0.0007** |
| | Day 14 vs. Day 28 | t(35)=2.447; p=0.0575 |
| | **Day 14 vs. Day 56** | **t(35)=2.724; p=0.0393** |
| | Day 28 vs. Day 56 | t(35)=0.4586; p=0.6494 |
| | **TBI** | |
| | Day 7 vs. Day 14 | t(35)=0.1231; p=0.9205 |
| | Day 7 vs. Day 28 | t(35)=1.253; p=0.6784 |
| | Day 7 vs. Day 56 | t(35)=1.432; p=0.6513 |
| | Day 14 vs. Day 28 | t(35)=1.094; p=0.6784 |
| | Day 14 vs. Day 56 | t(35)=1.297; p=0.6784 |
| | Day 28 vs. Day 56 | t(35)=0.3641; p=0.9205 |

**Table S9.** Statistical analyses related to Fig. 11: Wallerian degeneration is not required for homotopic collateral sprouting. Comparison of CTB(+) terminal densities in the superior colliculus of male and female, wt and *Sarm1* KO mice.

| **Two-way ANOVA** | **Post-hoc comparisons (WT vs *Sarm1* KO)** | |
|---|---|---|
| **Male** | | |
| Genotype<br>F (1, 57) = 0.2695; p=0.6056<br>Injury groups<br>F (3, 57) = 4.708; p=0.0053<br>Interaction<br>F (3, 57) = 0.4418; p=0.7240 | Sham | t(57)=0.1633; p=0.8889 |
| | D7 | t(57)=0.628; p=0.8889 |
| | D14 | t(57)=0.6486; p=0.8889 |
| | D56 | t(57)=0.8276; p=0.8799 |
| **Female** | | |
| Genotype<br>F (3, 51) = 2.241; p=0.0946<br>Injury groups<br>F (3, 51) = 4.418; p=0.0078<br>Interaction<br>F (3, 51) = 2.241; p=0.0946 | Sham | t(51)=0.5602; p=0.8217 |
| | D7 | t(51)=2.134; p=0.1315 |
| | D14 | t(51)=2.171; p=0.1315 |
| | D56 | t(51)=0.2125; p=0.8326 |
